# Rapid protamine evolution suppresses meiotic drive in *Drosophila*

**DOI:** 10.1101/2025.09.03.674087

**Authors:** Ching-Ho Chang, Aida Flor de la Cruz, Hung-Yu Lai, Isabel Mejia Natividad, Alex Noyola, Elian Angelo Magsino Abellanosa, Nicolas Lee, Harmit S. Malik

**Affiliations:** Division of Basic Sciences, Fred Hutchinson Cancer Center, United States; Institute of Molecular Biology, Academia Sinica, Taipei, Taiwan; Molecular and Cell Biology, Taiwan International Graduate Program, Academia Sinica; Graduate Institute of Life Sciences, National Defense Medical Center, Taipei, Taiwan; Howard Hughes Medical Institute, Fred Hutchinson Cancer Center, United States

## Abstract

Many animal species replace histones with protamines during spermatogenesis. Although essential for sperm function, protamines evolve rapidly for reasons that remain unknown. Using *in vivo* gene replacement, we examined the causes and consequences of the rapid evolution of *Mst77F*, a protamine essential for male fertility in *D. melanogaster*. Replacing *Mst77F* with divergent orthologs caused DNA compaction defects in X-bearing sperm, resulting in fewer mature X-bearing sperm and male-biased progeny. Reducing *D. melanogaster Mst77F* dosage caused the same bias, whereas increasing the dosage of *Mst77F* orthologs rescued it. Engineered sex-chromosome fusions revealed that *Mst77F* protects the *D. melanogaster* X chromosome from a killer-target meiotic drive system in a dosage- and sequence-dependent manner. Unlike in *D. melanogaster*, *Mst77F* was dispensable for fertility in *D. yakuba,* but remained important for suppressing sex-ratio distortion. Our findings provide the first evidence that relentless pressure to suppress sex-chromosome meiotic drive underlies the rapid evolution of protamines.

**One-sentence summary:** A rapidly evolving essential protamine suppresses sex-chromosome meiotic drive in *Drosophila*

## Main Text

Most eukaryotes rely on histones and histone variants to package their DNA and regulate processes such as transcription, DNA repair, and chromosome segregation (*1*). However, various animal species have independently evolved distinct DNA-compaction proteins, collectively referred to as protamines, to facilitate the compact packaging of genomes by up to 100-fold within sperm heads relative to somatic nuclei (*2–5*). For example, mammalian protamines are likely derived from linker histones (*6, 7*), whereas *Drosophila* protamines (or sperm nuclear basic proteins, SNBPs) are derived from HMG-box encoding transcription factors (*2, 8*). Although protamines are absent in some animals, they are essential for male fertility in all studied species that encode them, including mammals and *Drosophila* (*4*).

Despite their critical roles in sperm genome packaging and male fertility, protamine genes are among the fastest-evolving genes across various animal species, including mammals (*9–11*) and *Drosophila* species (*2*). A prevailing hypothesis suggests that the rapid evolution of protamines results from their role in shaping the sperm head and enhancing sperm success amid competition for fertilization (*10, 12, 13*). Indeed, protamine loss or mutations impair sperm head morphology and fertilization (*14–16*). However, there is no direct evidence that rapid protamine evolution is driven by a selective advantage for successful fertilization (*17*). Moreover, the intensity of sperm competition is either poorly correlated or not correlated with protamine evolutionary rates (*9, 10, 18, 19*). Some studies have alternatively suggested that the rapid evolution of protamines in mammals results from hypermutation at their CpG-bearing arginine codons, followed by selection to restore optimal arginine content for protamine function (*20*).

Our recent phylogenomic analysis of protamine genes in *Drosophila* species, which encode at least 15 distinct protamine genes (*21–24*), proposed an alternative explanation for the rapid evolution of protamines (*2*). Our analysis revealed extensive turnover of protamine genes associated with sex chromosome evolution, including multiple independent protamine gene duplications and amplification events on sex chromosomes across *Drosophila* species (*2, 25–28*). Such gene duplicates, including X-linked paralogs of *ProtA*/*B* and Y-linked paralogs of *Mst77F (Mst77Y)*, could evolve roles as sperm killers – selfish genes that enhance their propagation at the expense of host fertility (*25, 26, 29*). Conversely, in the *montium* subgroup of *Drosophila* species, half of the repertoire of ancestrally conserved protamine genes on autosomes was lost coincident with an X^Y chromosomal fusion, which obviated sex chromosome competition (*2*). These findings suggested that rapid protamine evolution may be driven by the need to suppress sex-chromosome meiotic drive, *i.e.,* the non-Mendelian over-transmission of one sex chromosome arising from conflict between X- and Y-bearing sperm, thereby maintaining male fertility and preserving the Fisherian optimal 50:50 sex ratio in resulting progeny (*30, 31*). However, the biological causes underlying the rapid evolution of protamines remained experimentally unverified in any species.

Here, using gene knockouts, ortholog replacements, and engineered sex-chromosome fusions, we show that the autosomal protamine *Mst77F* suppresses sex-chromosome meiotic drive in *D. melanogaster* and that its correct sequence and dosage are required for this suppression. This distortion arises from unequal maturation rates of sperm carrying X or Y chromosomes during spermatogenesis. In *D. melanogaster,* transgenic flies with reduced *Mst77F* dosage or orthologs from other species exhibit chromatin condensation defects in sperm bearing X chromosomes, resulting in their underrepresentation in mature sperm and ultimately leading to male-skewed progeny. Using engineered sex-chromosome fusions, we show that the distortion reflects a killer-target mechanism acting on X-linked sequences in *D. melanogaster*. Extending these findings to a second species, we show that knocking out *Mst77F* in *D. yakuba* does not cause complete sterility but does cause significantly male-biased progeny. Our results suggest that the need to suppress competitive interactions between sperm with different genotypes, such as X- and Y-bearing sperm, during spermatogenesis may be one of the key forces driving the rapid evolution of protamines in *Drosophila* and likely other animal species.

## Result

### *Mst77F* protamine replacements incompletely rescue male fertility in *D. melanogaster*

Using CRISPR/Cas9 and two guide RNAs (gRNAs), we generated a complete *Mst77F* knockout (*Mst77F KO*) by replacing all protein-coding sequences with an attP site, enabling subsequent site-specific ΦC31 recombinase-mediated insertion (Fig. 1A; Fig. S1A–B). Recapitulating earlier findings (*21, 32–34*), we observed that *Mst77F KO* homozygous males are entirely sterile, producing no mature sperm or offspring (Fig. 1B; Fig. S1C). Spermatids in *Mst77F KO* males still appear to undergo the histone-to-protamine transition, removing most histones and incorporating other protamines, *e.g., ProtA/B* (*34*). However, we observed that some spermatids have larger nuclei (Fig. S1D), consistent with *Mst77F*’s proposed role in condensing genomic DNA in sperm.

**Fig. 1:**
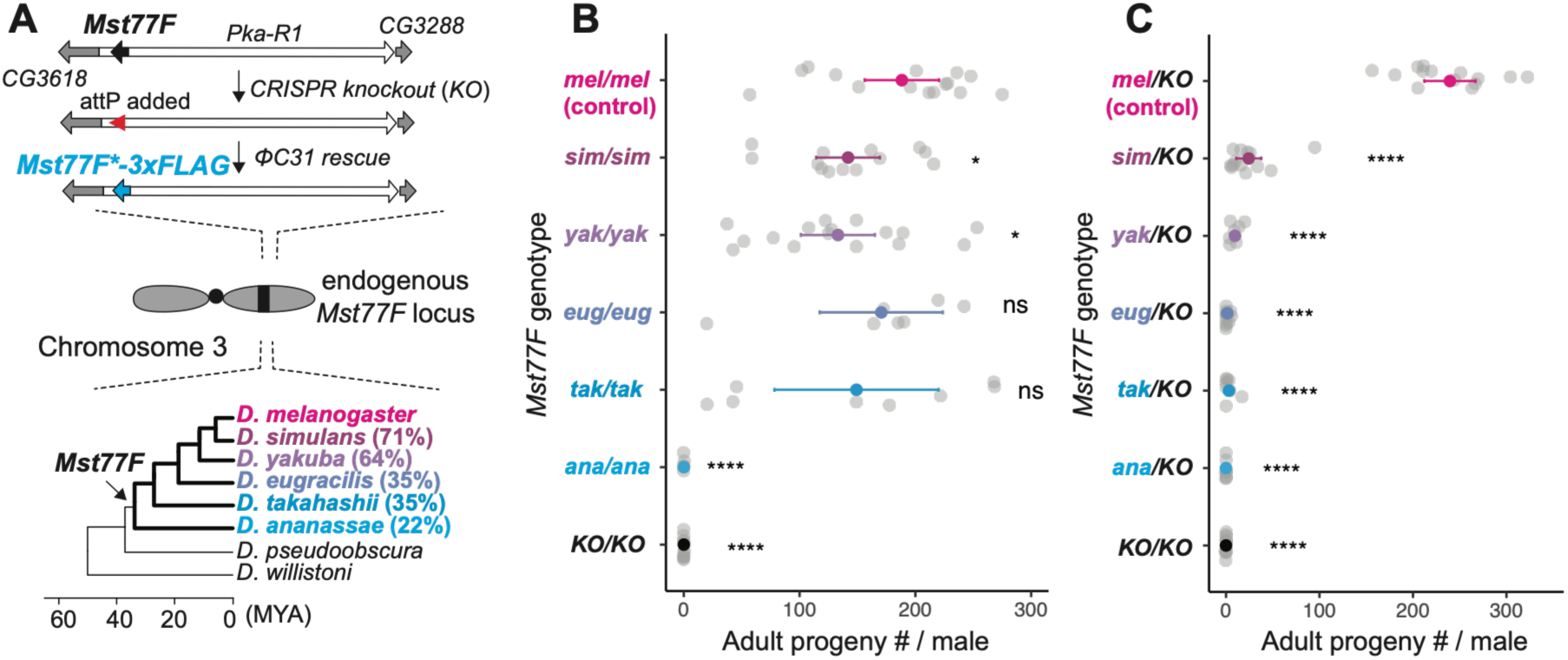
Functional divergence of the *Mst77F* essential protamine across *Drosophila* species. **(A)** Schematic of the CRISPR/Cas9-mediated knockout and ΦC31 integrase-mediated replacement of *Mst77F* in *D. melanogaster*. The endogenous *Mst77F* is located within an intron of *Pka-R1*. It was first replaced with a *DsRed* marker (not shown) and an attP site, generating male-sterile *Mst77F* knockouts (*KO*). Orthologous *Mst77F* transgenes from various *Drosophila* species were then inserted into the introduced attP site and tested for their ability to rescue male fertility. A phylogeny shows divergence of *Drosophila* species from *D. melanogaster* (millions of years ago, MYA). Also shown is the % protein identity between each ortholog and Mst77F-mel. **(B)** Comparison of the fertility of transgenic homozygous *D. melanogaster* males carrying two copies of *Mst77F* from different species. Each dot represents the total number of adult offspring produced by a single male. (**C**) Comparison of the fertility of transgenic hemizygous *D. melanogaster* males encoding only a single copy of *Mst77F* from the designated *Drosophila* species. Statistics for progeny counts were carried out using unpaired Student’s *t*-tests (*p<0.05; ****p < 0.0001; ns, not significant).

To rescue the male sterility caused by the loss of *Mst77F*, we cloned and reintroduced the wild-type *D. melanogaster* cDNA copy of *Mst77F* (*Mst77F-mel*) using ΦC31-mediated integration at the endogenous locus (Fig. 1A). The construct included 500 bp of endogenous upstream sequence as the promoter and carried a C-terminal 3×FLAG epitope tag. As previously reported (*21, 32–34*), either one or two copies of *Mst77F-mel* in the endogenous locus could rescue the *Mst77F* knockout, with each male capable of producing more than 100 offspring (Fig. 1B-C; Fig S1C; Data S2). This shows that loss of *Mst77F* causes male sterility and that the 3×FLAG tag does not interfere with its essential function in male fertility. Immunostaining with a FLAG antibody revealed that transgenic Mst77F is incorporated into spermatid chromatin during the late canoe stage of spermatogenesis (Fig. S2), like endogenous Mst77F (*34*).

Based on phylogenetic evidence, we infer that *Mst77F* is one of the youngest *Drosophila* protamine genes, likely originating in the common ancestor of *D. melanogaster* and *D. ananassae* ∼20 million years ago (Fig. 1A) (*35*). If the rapid evolution of *Mst77F* were tied to its roles in male fertility, we hypothesized that orthologs from various species might be maladapted for one or more of these roles in *D. melanogaster*, potentially revealing the selective pressures driving *Mst77F’s* rapid divergence. Therefore, we utilized the attP site introduced at the endogenous *Mst77F* locus to introduce an allelic series of C-terminal 3×FLAG-tagged *Mst77F* orthologs from five divergent *Drosophila* species encoding proteins with increasing amino acid divergence (Fig. 1A; Data S1): *D. simulans* (2.5 mya, *Mst77F-sim,* 71% amino acid identity)*, D. yakuba* (6 mya, *Mst77F-yak,* 64%)*, D. eugracilis* (10 mya, *Mst77F-eug,* 35%)*, D. takahashii* (15 mya, *Mst77F-tak,* 35%), and *D. ananassae* (25mya, *Mst77F-ana,* 22%). We only swapped the protein-coding sequences, retaining the regulatory sequences from the *D. melanogaster Mst77F* locus. We confirmed that all *Mst77F* transgenes were expressed at similar stages of spermatogenesis and comparable levels to the FLAG-tagged *Mst77F-mel* transgene using immunostaining (Fig. S2A) and transcriptomic analyses (Fig. S3A). Subsequent Western blot analyses (Fig. S3B-C; Data S3), with or without Calf Intestinal Alkaline Phosphatase (CIP) treatment, revealed that all transgenic Mst77 proteins, except Mst77F-ana, are phosphorylated in *D. melanogaster* (Fig. S3B). Consistent with normal completion of the histone-to-protamine transition, we confirmed that the histone variant His2Av is correctly removed in sperm from transgenic flies, whereas the centromeric histone Cid is retained (Fig. S2B), as previously reported (*36*).

We first assessed whether *Mst77F* ortholog replacements affected male fertility in *D. melanogaster.* We found that *D. melanogaster* males homozygous for *Mst77F-ana* were sterile, like the *Mst77F-mel* knockout (Fig. 1B; Fig. S1C). In contrast, transgenic males homozygous for other *Mst77F* orthologs had only a modest decrease in fertility at 25°C (Fig. 1B) or at 29°C (Fig. S4A), a temperature known to perturb sperm development in *Drosophila* (*37*). However, we observed a slight but consistent decrease in seminal vesicle size among transgenic males homozygous for heterospecific *Mst77F* orthologs compared to *Mst77F-mel* homozygous males (Fig. S2B; Data S4). Thus, our findings show that most homozygous *Mst77F* orthologs from closely related species can complement the crucial male fertility function in *D. melanogaster*, despite sharing only 35–71% protein identity. Only Mst77F-ana, which is 22% identical to Mst77F-mel, cannot. Since Mst77-ana is also the only Mst77F protein that is not phosphorylated in *D. melanogaster*, our findings suggest that phosphorylation may be a critical post-translational requirement for *Mst77F* to fulfill its essential role in spermiogenesis (*32*).

We next reduced *Mst77F* dosage by generating hemizygous males with one *Mst77F* allele and one knockout allele. Whereas hemizygous (single-copy) *Mst77F-mel* could restore wild-type male fertility, the hemizygous *Mst77F-sim*, *Mst77F-yak*, *Mst77F-eug,* and *Mst77F-tak D. melanogaster* strains showed more than a ten-fold reduction in fertility (Fig. 1C). Thus, *Mst77F* orthologs from closely related species are haploinsufficient for normal male fertility in *D. melanogaster*, but this impairment can be relieved by increased dosage (Fig. 1B).

### *Mst77F* protamine replacements unleash sex-ratio distortion in *D. melanogaster*

We previously hypothesized that autosome-encoded protamines, such as *Mst77F,* might evolve rapidly to suppress competition between X- and Y-bearing sperm, thereby ensuring optimal sex ratios and high male fertility. To test this hypothesis, we evaluated how the *Mst77F* gene replacements affected the sex ratio of the resulting adult progeny. We crossed transgenic *D. melanogaster* (XY) males encoding two copies of *Mst77F-mel* to wild-type *D. melanogaster* females (Fig. 2A). These crosses produced ∼50% males at both 25°C (Fig. 2B–C) and 29°C (Fig. S4B), consistent with Mendelian expectations, indicating equal success of X- and Y-bearing sperm.

**Fig. 2:**
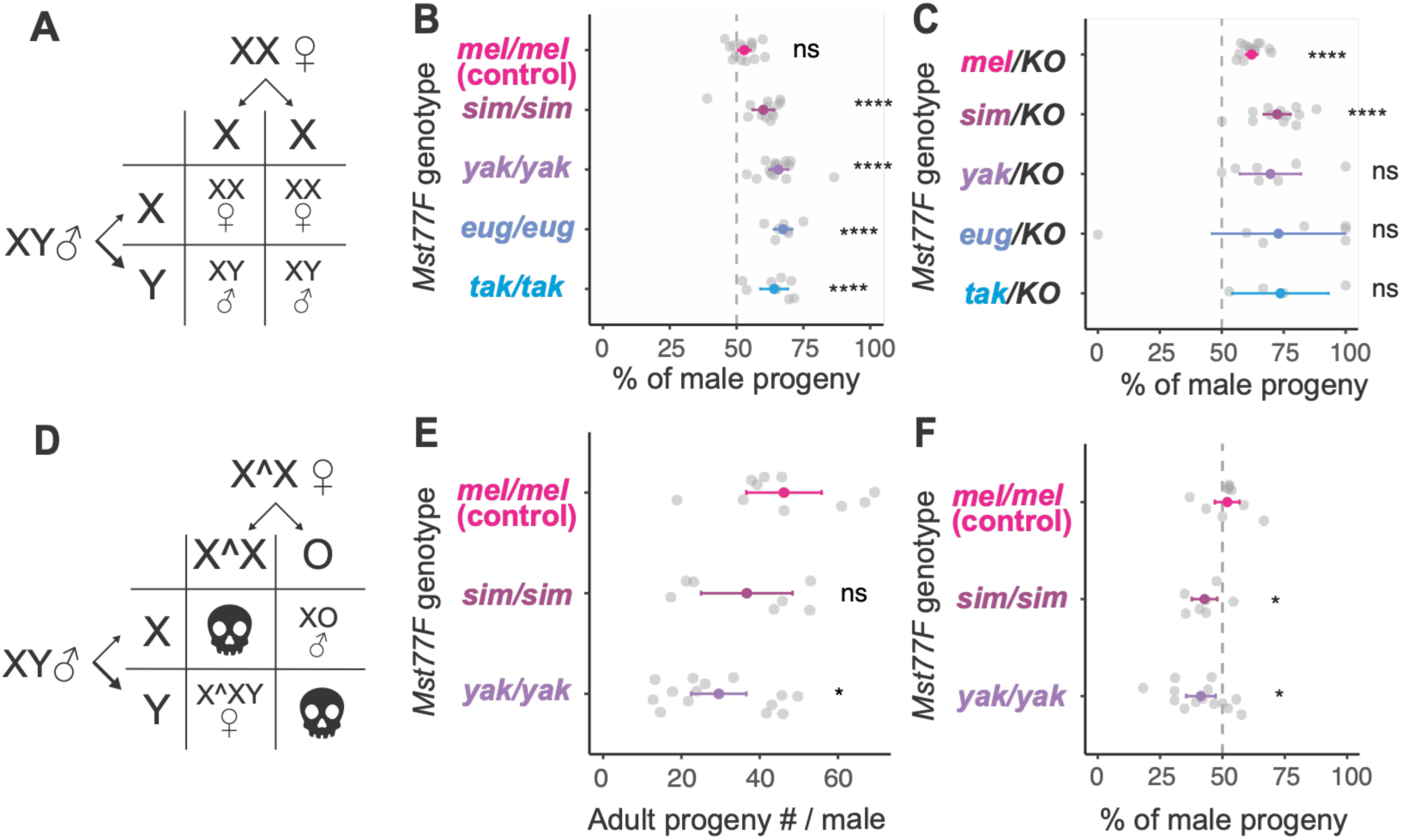
*Mst77F* gene replacements in *D. melanogaster* generate progeny with a biased sex ratio. (**A**) Punnett squares illustrate the expected genotypes from crosses between transgenic *Mst77F*-encoding XY males and wild-type XX females. Sex ratios of resulting adult progeny from transgenic *D. melanogaster* males carrying (**B**) two copies of *Mst77F* from the designated species, or (**C**) one copy of *Mst77F* from the designated species. Each dot represents data from a single male (n ≥ 7). The dashed line indicates the expected 50:50 Mendelian ratio. In some cases, low fertility (<100 offspring/male) of hemizygous males precludes robust tests against a standard 50:50 ratio using χ² test. (**D**) Punnett square illustrates the expected genotypes from crosses between transgenic *Mst77F*-encoding XY males and females carrying an attached-X chromosome (X^X). In matings with X^X females, fertilization by X-bearing sperm produces XO males, whereas fertilization by Y-bearing sperm produces X^XY females. Embryos carrying zero or three X chromosomes are inviable. (**E**) In crosses with X^X females, transgenic homozygous *D. melanogaster* males carrying two copies of *Mst77F-mel* have higher fertility than *D. melanogaster* males carrying two copies of *Mst77F-sim* or *Mst77F-yak.* (**F**) Transgenic homozygous *D. melanogaster* males carrying two copies of *Mst77F-mel* produce progeny with expected 50:50 Mendelian ratios. In contrast, *D. melanogaster* males carrying two copies of *Mst77F-sim* or *Mst77F-yak* produce more female-biased progeny. Statistics for deviations from the expected 50% male progeny ratio were carried out using χ² tests and statistical comparisons of transgenic flies to transgenic flies encoding homozygous *Mst77F- mel* (control) using unpaired Student’s *t*-tests (*p<0.05; **p<0.01; ***p<0.001; ****p < 0.0001; ns, not significant).

In contrast, fertile males with two copies of *Mst77F* orthologs, even from closely related species, produced over 60% male offspring. The sex-ratio distortion was further exacerbated by halving the dosage of *Mst77F*. Indeed, even *D. melanogaster* males carrying a single copy of *Mst77F-mel* produced 62% male progeny (Fig. 2C) even though these males appear to be highly fecund (Fig. 1C). Males carrying only a single copy of *Mst77F* orthologs from other species yielded approximately 70% male progeny (Fig. 2C). Conversely, both the fertility defects and the sex-ratio distortion were alleviated by increasing the dosage of heterospecific *Mst77F-yak* (Fig. S5A-B). Moreover, heterozygous males carrying one copy of *Mst77F-mel* along with a single copy of an *Mst77F* ortholog performed better than *Mst77F-mel* hemizygous males. Thus, unlike *Mst77F-mel,* heterospecific *Mst77F* orthologs cannot fully rescue male fertility or suppress sex-ratio distortion, but they do not act as dominant negatives in *D. melanogaster* (Fig. S5C–D). These findings suggest that both protein divergence and gene dosage of the *Mst77F* orthologs affect their ability to suppress sex-ratio distortion.

Sex-ratio distortion can arise from differential production or fertilization success of X- or Y-bearing sperm, or from viability differences between female or male offspring. To differentiate between these possibilities, we crossed *Mst77F-mel*, *Mst77F-sim*, or *Mst77F-yak* homozygous male (XY) flies with females possessing attached-X (or X^X) chromosomes (Fig. 2D). In this cross, viable progeny can only result from nullo-X eggs fertilized by X-bearing sperm, which develop into XO males, or X^X eggs fertilized by Y-bearing sperm, which develop into X^XY females (*Drosophila* sex is determined by the X: autosome ratio, rather than the Y chromosome). This system effectively reverses the inheritance patterns of the X and Y chromosomes, allowing us to resolve confounding factors such as post-zygotic female-specific lethality. We observed slightly lower fertility among transgenic males homozygous for heterospecific *Mst77F* orthologs compared to *Mst77F-mel* homozygous males (Fig. 2E). We found no skew in progeny sex ratios in crosses between *Mst77F-mel* homozygous males and X^X females (Fig. 2F). In contrast, we noted a higher proportion of X^XY females among the progeny from crosses of *Mst77F-sim* or *Mst77F-yak* homozygous males with X^X females compared to *Mst77F-mel* homozygous males (Fig. 2F). Thus, both crosses to XX and X^X females produced fewer progeny derived from X-bearing sperm. Because X-bearing sperm give rise to female progeny in the standard cross but to XO male progeny in the attached-X cross, this consistent deficit cannot be explained by sex-specific offspring viability. Instead, an overrepresentation of Y-bearing sperm relative to X-bearing sperm must underlie this skew, occurring either during spermatogenesis or at fertilization.

To test whether suppression of sex-chromosome meiotic drive is a general property of *Drosophila* protamines or specific to *Mst77F*, we examined flies deleted for the protamines *ProtA* and *ProtB*, which have previously been implicated in suppressing the autosomal meiotic driver *Segregation Distorter* (*SD*) (*38*). Unlike perturbations of *Mst77F*, deletion of *ProtA* and *ProtB* did not skew progeny sex ratios (n=2918 from 10 individual males; χ² test, P>0.9; Data S2). This indicates that suppression of sex-chromosome drive is not shared by all protamines, but it raises the possibility that different protamines may have evolved to counteract distinct meiotic drivers.

### Multiple domains of *Mst77F* contribute to its fertility and drive-suppression functions

*Mst77F-yak* is impaired for both fertility-essential and meiotic drive suppressor functions in *D. melanogaster* (Fig. 3A), even though it is separated from *Mst77F-mel* by merely six million years. Both Mst77F-mel and Mst77F-yak proteins are of similar length (215 and 219 amino acid residues, respectively) but differ by 78 amino acids, highlighting the remarkably rapid sequence evolution of *Mst77F*. These amino acid differences are distributed throughout the proteins, except for the relatively conserved high mobility group (HMG) domains and predicted nuclear localization signals (NLS; Fig. 3A). To identify the domains associated with fertility-essential or meiotic drive suppressor functions, we constructed chimeric proteins comprising different combinations of four segments of *Mst77F-mel* and *Mst77F-yak* (segments numbered 1–4 from N- to C-terminus; Fig. 3A).

**Fig. 3:**
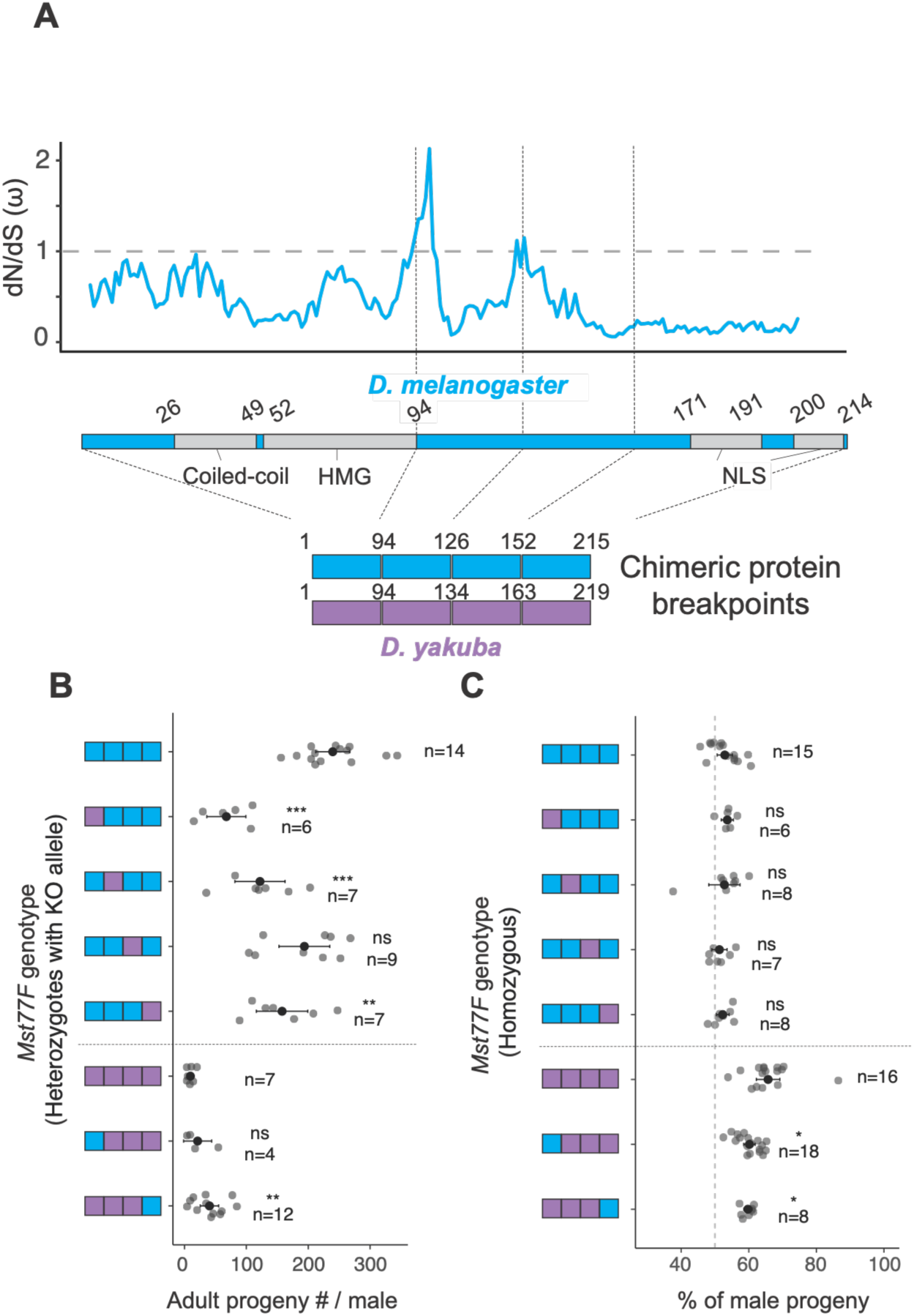
Fertility and progeny sex ratios of *D. melanogaster*-*D. yakuba Mst77F* chimeras. **(A)** Schematic representation of predicted protein domain structures of Mst77F in *D. melanogaster* (*2, 49*), including a coiled-coil domain (aa 26-49), a high mobility group (HMG) box DNA-binding domain (aa 52-94), and two putative nuclear localization signals (NLS, aa 171-191, 200-214). The top panel shows a sliding-window analysis of dN/dS (ω) across the Mst77F coding sequence, highlighting a peak of positive selection (dN/dS > 1) just downstream of the HMG-box. Numbers indicate the positions of three breakpoints used to generate chimeric proteins for *D. melanogaster* and *D. yakuba*. Colors in the chimeric bars represent the species’ origin of the Mst77F segment, with *D. melanogaster* in blue and *D. yakuba* in magenta. **(B)** We measured the fertility of hemizygous *D. melanogaster* transgenic males expressing a single copy of *Mst77F-mel*, *Mst77F-yak*, or chimeric genes. Each dot represents data from an individual male (the number of males tested is also reported). **(C)** We measured the adult progeny sex ratio of *D. melanogaster* transgenic males expressing two copies of *Mst77F-mel*, *Mst77F-yak*, or chimeric genes. The number of males tested is also reported. Statistics for deviations from the expected 50% male progeny ratio were carried out using χ² tests and statistical comparisons of transgenic flies encoding chimeric genes were carried out to transgenic flies encoding either *Mst77F- mel* (above dashed line) or *Mst77F-yak* (below dashed line) using unpaired Student’s *t*-tests (*p<0.05; **p<0.01; ***p<0.001; ****p < 0.0001; ns, not significant).

Single hemizygous *Mst77F-mel* males produced more than 200 offspring on average, whereas hemizygous *Mst77F-yak* males produced fewer than 20 progeny (Fig. 1). Therefore, to provide the most sensitive assay for male fertility, we tested each chimeric *Mst77F* gene in a single copy. Swapping the first, second, or fourth segment from *Mst77F-yak* into *Mst77F-mel* resulted in a significant reduction in fertility, whereas swapping in the third segment did not impact fertility. We also conducted a reciprocal analysis and discovered that the first or fourth segment from *Mst77F-mel* could only marginally rescue the low fertility of *Mst77F-yak* males (Fig. 3B). Hence, optimal fertility requires multiple domains of *Mst77F-mel* (Fig. 3B).

We next investigated the *Mst77F* chimeras for their ability to suppress sex-ratio distortion. *Mst77F-mel* homozygous males show no sex-ratio distortion, whereas *Mst77F-yak* homozygous males yield approximately 65% male progeny (Fig. 3C). Thus, to enhance the sensitivity of our comparison, we tested all chimeras as homozygotes, which also provided the added benefit of relatively higher overall fertility. We found that no single segment from *Mst77F-yak* significantly impaired *Mst77F-mel*’s capacity to suppress sex-ratio distortion (Fig. 3C).

Conversely, introducing either the first or the fourth segment from *Mst77F-mel* into *Mst77F-yak* significantly alleviated but did not eliminate sex-ratio distortion (Fig. 3C). These results confirm that multiple segments of *Mst77F-mel* contribute to suppressing sex-ratio distortion.

By comparing all seven *Mst77F* transgenes, including *Mst77F-mel*, *Mst77F-yak*, and their chimeras (n=7), we identified a strong anti-correlation between male fertility and sex-ratio distortion (Pearson’s R=-0.82, p=0.01; Spearman’s rank correlation ρ = -0.857, p=0.01). This anti-correlation suggests that the depletion of X-bearing sperm contributes to both sex-ratio distortion and decreased male fertility.

### X-chromosome-specific DNA condensation defects in spermatids underlie the sex-ratio distortion in *Mst77F* replacement males

To investigate whether X- and Y-bearing gametes progress similarly through spermatogenesis, we used fluorescent *in situ* hybridization (FISH) to detect satellite DNA located on either Y chromosomes (*AATAC*) or autosomes (*Prod-sat*) (Fig. 4A). This strategy allowed us to distinguish X-bearing mature sperm, which hybridize solely with the *Prod-sat* probe, from Y-bearing mature sperm, which hybridize with both the *Prod-sat* and *AATAC* probes. We then calculated the proportions of X- and Y-bearing mature sperm in transgenic *D. melanogaster* males encoding two copies of different *Mst77F* orthologs. We found no X-versus-Y skew in mature sperm from *Mst77F-mel* homozygous males (Fig. 4B–C), consistent with no observed sex-ratio distortion in progeny from these males (Fig. 2). In contrast, we identified a significant overrepresentation of Y-bearing sperm in hemizygous *Mst77F-mel* males encoding a single copy of *Mst77F-mel,* or in *Mst77F-sim, Mst77F-yak, Mst77F-eug,* or *Mst77F-tak* homozygous males (Fig. 4B–C; Fig. S6A; Data S5), corresponding with the sex-ratio bias observed in their resulting adult progeny (Fig. 2). These findings suggest that the under-representation of X-bearing sperm during spermatogenesis is sufficient to explain the sex-ratio distortion observed in the progeny of *Mst77F* replacement males.

**Fig. 4:**
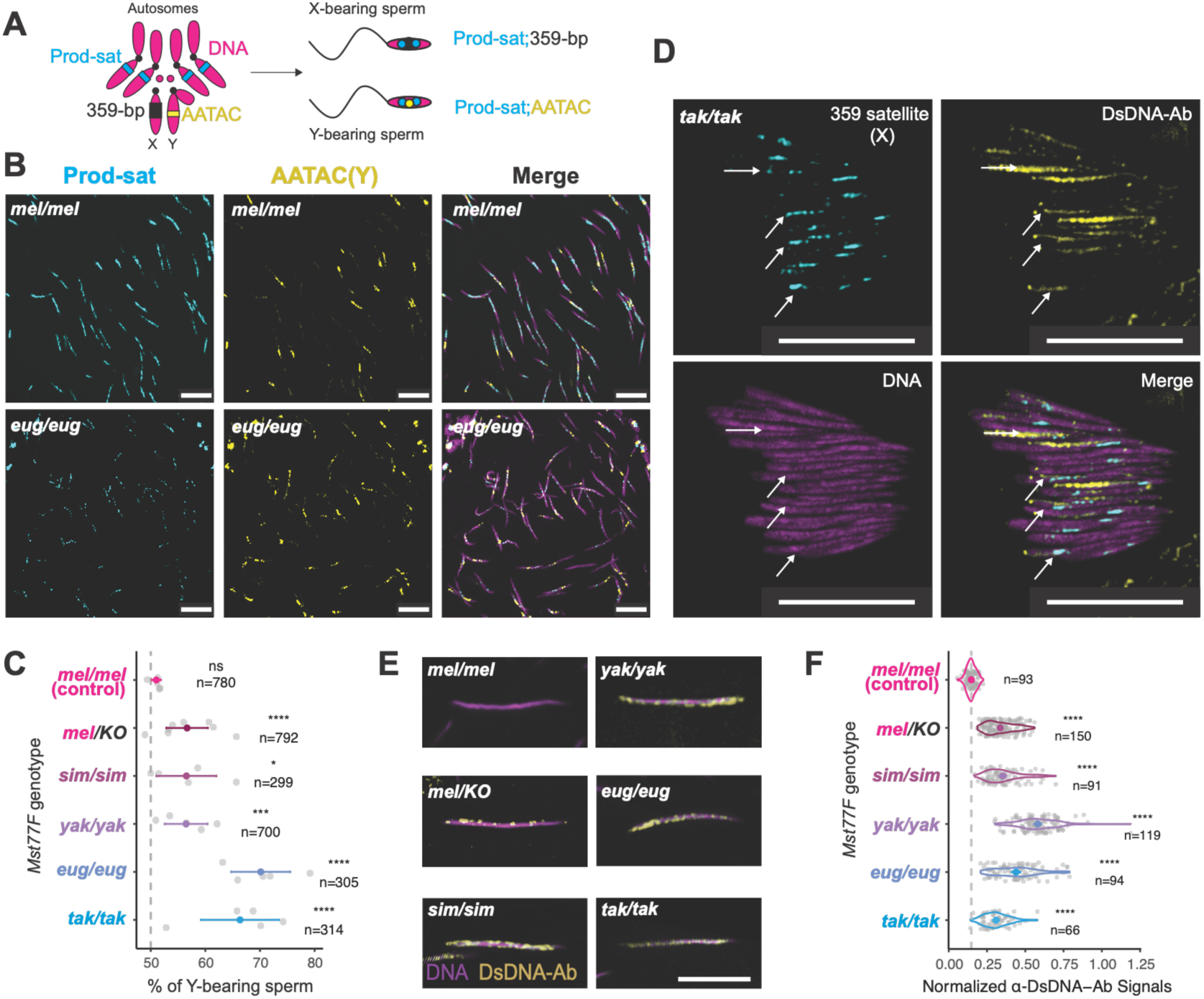
*Mst77F* gene replacements preferentially disrupt sperm chromatin condensation and decrease the proportion of X-bearing sperm. (**A**) Schematic of DNA FISH probes used to distinguish X- and Y-bearing sperm based on the Y-linked AATAC (yellow), X-linked 359-bp satellite (black), and autosomal Prod-sat (cyan) satellite repeats. Y-bearing spermatids and sperm hybridize to both AATAC and Prod-sat probes, while X-bearing spermatids and sperm hybridize to Prod-sat and 359-bp probes. DNA was stained with Hoechst 33342 (magenta). (**B**) FISH was performed using the Y-linked AATAC (yellow) and Prod-sat (cyan) probes on seminal vesicles from homozygous *D. melanogaster* males encoding either two copies of *Mst77F-mel* or two copies of *Mst77F-eug*. The results from other genotypes are shown in Fig. S6A. (**C**) Quantification of the proportion of Y-bearing sperm among total mature sperm in each of the six transgenic *Mst77F* gene replacement genotypes from (B and S6A). Each dot represents data from one seminal vesicle, and the dashed line represents the expected 50:50 Mendelian ratio. (n = total number of surveyed sperm in each genotype; χ² test against 50:50 ratio). (**D**) Immunostaining using an α-dsDNA antibody (yellow) was combined with FISH analyses using a probe against X-linked 359-bp satellite repeats (cyan) on late-stage spermatids from *Mst77F*-tak replacement males. DNA was stained with Hoechst 33342(magenta). White arrows highlight spermatids with signals from both the X-linked 359 probe and α-dsDNA antibody. The results from other genotypes are shown in Fig. S6B. **(E)** Mature sperm were stained with an α-dsDNA antibody (yellow) to assess their degree of chromatin compaction. Representative images of single sperm are shown here; seminal vesicle images are shown in Fig. S6C. **(F)** α-DsDNA antibody staining was carried out on sperm from the various *Mst77F* replacement *D. melanogaster* males. Images were quantified and normalized to the Hoechst 33342 signal. A minimum of 60 nuclei were assayed for each genotype. Statistical analyses were performed using unpaired Student’s *t*-tests (*p<0.05; ***p<0.001; ****p < 0.0001; ns, not significant). Scale bar = 10 μm for all images.

Since *Mst77F* is critical for DNA condensation, we hypothesized that delays or defects in DNA condensation might underlie reduced fertility and sex-ratio distortion in males with improper *Mst77F* dosage or sequence. Under this hypothesis, X-bearing spermatids are more prone to DNA condensation defects than Y-bearing spermatids. To test this possibility, we focused on *Mst77F-tak* homozygous *D. melanogaster* flies, which have the most severe male bias in their progeny (Fig. 2B). In *Drosophila* species, X- and Y-bearing sperm resulting from a single meiotic event are produced simultaneously and undergo synchronous replication, resulting in a cyst containing 32 X-bearing and 32 Y-bearing spermatids. We performed immunostaining of spermatid cysts from *Mst77F* replacement homozygous *D. melanogaster* flies using an antibody to double-stranded DNA (α-dsDNA), which identifies decondensed spermatids (*39*). Under normal circumstances, chromatin in late spermatids and mature sperm is so tightly compacted that it completely excludes the α-dsDNA antibody. However, DNA condensation defects would result in staining with the α-dsDNA antibody (Fig. 4D–F; Fig. S6B–C). To distinguish whether spermatids containing X- or Y-chromosomes might be more prone to condensation defects, we combined immunostaining using the α-dsDNA antibody with FISH using an X-chromosome-specific *359-bp* probe (Fig. 4A) in *Mst77F-tak* homozygous *D. melanogaster* flies. We found that, prior to spermatid individualization, ∼84% of decondensed spermatids are X-bearing (n=57; p<0.001; Fig. 4D). In some cysts, well-condensed Y-bearing spermatids appear to individualize earlier than neighboring X-bearing spermatids, which remain poorly condensed. Thus, in *Mst77F-tak* homozygous males, decondensing spermatids are predominantly X-bearing. We infer this is also the case for the other *Mst77F* ortholog replacement males based on their mature-sperm ratios (Fig. 4B–C) and progeny sex ratios (Fig. 2).

We investigated whether chromatin condensation defects persist in mature sperm, even after sperm individualization. We observed no or minimal α-dsDNA antibody staining in mature sperm from *Mst77F-mel* homozygous *D. melanogaster* males (Fig. 4E–F; Fig. S6C). In contrast, we observed significant α-dsDNA antibody staining in mature sperm either from hemizygous *Mst77F-mel* males or from homozygous *Mst77F-sim, Mst77F-yak, Mst77F-eug,* or *Mst77F-tak* males, consistent with defective DNA condensation persisting even in mature sperm (Fig. S6C). We conclude that the inadequate dosage of conspecific *Mst77F-mel* or the inappropriate sequence of heterospecific *Mst77F* leads to defective chromatin condensation and differential loss of X-bearing sperm during sperm development in *D. melanogaster,* resulting in meiotic drive. These same defects likely also underlie the gross alterations in sperm nuclear morphology that we observed in *Mst77F-mel* hemizygous males and homozygous *Mst77F* ortholog-replacement males, but not in homozygous *Mst77F-mel* males (Fig. S7A).

### *D. melanogaster Mst77F* suppresses a ‘killer-target’ meiotic drive targeting an X-chromosomal locus

We considered two possibilities to explain the X-chromosome condensation defects we observed in cases of inappropriate *Mst77F* dosage and sequence. The first possibility is akin to a ‘toxin-antidote’ system. Under this model, the Y chromosome encodes both a trans-acting ‘toxin’ factor that induces widespread chromatin decondensation and a cis-acting protective ‘antidote’ factor that specifically protects Y-bearing sperm from decondensation (Fig. 5A, left). Such ‘toxin-antidote’ meiotic drive systems have been previously observed on several occasions across taxa (*40–44*). Under this model, Y-bearing sperm would sequester both the toxin and the antidote, whereas X-bearing sperm would only sequester the toxin. Under this model, two copies of *Mst77F-mel* in wild-type *D. melanogaster* males produce sufficient Mst77-mel to counteract the toxin, leading to equal success of X- and Y-bearing sperm (Fig. 5B, left). However, *D. melanogaster* males, which are either hemizygous for *Mst77F-mel* or homozygous for *Mst77F* orthologs, cannot sufficiently counteract the toxin, leading to impairment of X-bearing sperm and male-biased progeny (Fig. 5C, left).

**Fig. 5.**
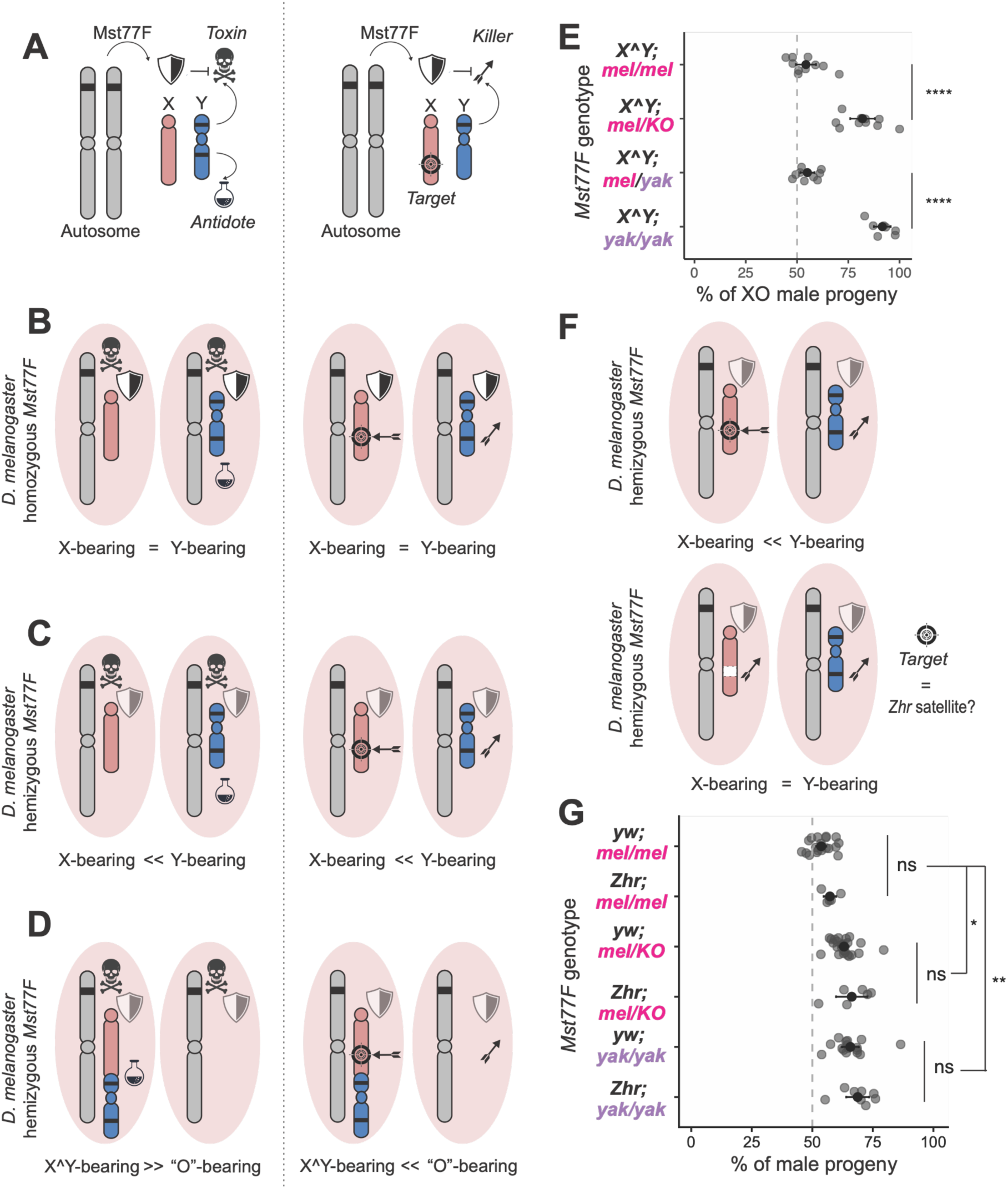
Genetic models of *Mst77F*-mediated sex-ratio distortion. **(A)** Alternative mechanistic models for the Mst77F-mediated meiotic drive. We propose that autosome-encoded Mst77F suppresses a trans-acting Y-linked toxin or killer which induces meiotic drive. In the Toxin-Antidote model (left), the Y chromosome, but not the X chromosome, encodes another cis-acting antidote to neutralize this trans-acting Y-linked toxin. In the Killer-Target model (right), the X chromosome carries a target that is susceptible to this Y-linked killer. When Mst77F functions properly, it suppresses the Y-linked toxin or killer, allowing sperm bearing either the X chromosome or Y chromosome to survive symmetrically **(B)**. When Mst77F function is disrupted—either by reducing gene dose (hemizygosity) or replacement by its ortholog—the activated Y-linked toxin or killer will eliminate sperm without the Y chromosome (Toxin-Antidote model) or carrying the X chromosome (Killer-Target model). In the case of males carrying independent X and Y chromosomes, the outcomes of both models are the same, leading to the overrepresentation of Y-bearing sperm **(C)**. To distinguish these two models, we disrupt *Mst77F* function in males carrying the fused X^Y chromosome, which produce sperm with either an X^Y chromosome or no sex chromosome (“O”-bearing). Under the Toxin-Antidote model, “O”-bearing sperm lack the Y-linked antidote and are eliminated. Under the Killer-Target model, the X^Y chromosome still retains the X-linked target and is eliminated **(D)**. **(E)** Dot plot illustrating the percentage of XO male progeny generated from O-bearing sperm across different Mst77F genetic backgrounds. Dashed line denotes the standard Mendelian expectation (50%). Statistical comparisons of flies were carried out to flies encoding either homozygous *Mst77F-mel* (mel/mel) or one copy of *Mst77F- mel* and *Mst77F-yak* (mel/yak) using unpaired Student’s *t*-tests (*p<0.05; **p<0.01; ***p<0.001; ****p < 0.0001; ns, not significant).

An alternative possibility is a ‘killer-target’ system, in which a trans-acting ‘killer’ factor acts on specific ‘target’ chromosomal regions to induce chromosome-specific decondensation (Fig. 5A, right). Such a ‘killer-target’ system is observed in sperm from *D. melanogaster* males that encode the *Segregation Distorter* (*SD*) autosomal meiotic driver, which is known to selectively kill sperm encoding large arrays of the *Responder* target satellite DNA on the 2^nd^ chromosome (*39*). Under this model, since the target of the drive is on the X chromosome, only X-bearing sperm will incur the consequences of the drive. Under this second model, two copies of *Mst77F-mel* produce sufficient Mst77-mel to counteract the ‘killer’ in wild-type *D. melanogaster* males, leading to equal success of X- and Y-bearing sperm (Fig. 5B, right). However, *D. melanogaster* males, which are either hemizygous for *Mst77F-mel* or homozygous for *Mst77F* orthologs, cannot sufficiently counteract the ‘killer’, leading to impairment of X-bearing sperm and male-biased progeny (Fig. 5C, right).

To distinguish between these two models, we used an attached XY (X^Y) chromosome, in which an X chromosome is fused to a Y chromosome, forming a single chromosome. X^Y males produce X^Y-bearing sperm and nullo-X sperm (lacking any sex chromosome) in equal proportions in the absence of any distortion. However, if distortion were to occur due to a toxin-antidote system, in which the Y chromosome encodes a cis-acting antidote, sperm carrying the X^Y chromosome would undergo normal sperm maturation despite the presence of X-linked sequences (Fig. 5D, left), whereas nullo-X sperm would sequester only the toxin and be impaired in chromatin condensation. This would lead to an over-representation of XX^Y females over XO males in the resulting progeny. Conversely, if distortion occurs due to a killer-target system that targets a locus on the X chromosome, X^Y-bearing sperm would bear the brunt of the decondensation and spermatogenesis defects, regardless of the attached Y-linked sequences (Fig. 5D, right), whereas nullo-X sperm would be unimpaired. This would lead to an excess of XO males over XX^Y females in the resulting progeny. Thus, the ratio of XX^Y females to XO males in the progeny of *Mst77F*-swap males can effectively discriminate between the two potential modes of distortion.

As a control, we first crossed wild-type *Mst77F-mel* X^Y males to females from the Oregon-R strain of *D. melanogaster*. Due to a lack of distortion, we observed a balanced 1:1 ratio of XX^Y female to XO male offspring, as expected (Fig. 5E). However, in crosses involving either hemizygous *Mst77F-mel* X^Y males or homozygous *Mst77F-yak* X^Y males, we observed a strongly distorted sex ratio of resulting progeny (>80% XO males) (Fig 5E; Data S2). These findings rule out the toxin-antidote model and strongly favor the killer-target model of distortion. They implicate a *cis*-acting sequence on the X chromosome as the target of distortion, leading to chromatin condensation failure and sperm maturation defects observed in *Mst77F*-swap males.

Given the gross condensation defects, we hypothesized that this *cis*-target sequence must be abundant and either unique to or significantly enriched on the *D. melanogaster* X chromosome. Based on these dual criteria, we investigated whether the X-linked target is the 1.688 satellite (also known as the 359 bp satellite or *Zhr,* for *Zygotic hybrid rescue*), a multi-megabase pericentromeric satellite DNA that recently expanded on the X chromosome of *D. melanogaster* (*45*). If *Zhr* represents the X-linked target of distortion, we predicted that an X chromosome carrying a large *Zhr* pericentromeric deletion (Δ*Zhr*) would be less susceptible to sex-ratio distortion caused by *Mst77F* mismatches (Fig. 5F). Contrary to this prediction, we found that the Δ*Zhr* X chromosome was not significantly less sensitive to sex-ratio distortion seen in wild-type *Mst77F-yak* homozygous *D. melanogaster* males (Fig. 5G). Even though our findings rule out *Zhr* as the target of drive, they indicate that an as-yet-unknown *D. melanogaster* X chromosomal locus is the target of the sex-chromosome meiotic drive unleashed in *Mst77F*-swap males.

### *D. yakuba Mst77F* knockouts are fertile but undergo sex-ratio distortion

The crucial function of *Mst77F* in male fertility is unlikely to be universally conserved across *Drosophila* species, given its recent emergence ∼20 million years ago and its loss in at least one lineage (the *montium* group). To clarify whether fertility, suppression of meiotic drive, or both are the primary selective pressures driving the rapid evolution of *Mst77F*, we generated a CRISPR-Cas9-mediated knockout of *Mst77F* in *D. yakuba,* a closely related species that diverged from *D. melanogaster* only 6 million years ago (Fig. S8)(*35, 46*). We discovered that, unlike in *D. melanogaster*, *Mst77F-yak* knockouts in *D. yakuba* exhibited reduced fertility but are not completely sterile (Fig. 6A), suggesting that *Mst77F*’s essential role in male fertility is likely a recently derived function in *D. melanogaster*.

**Fig. 6:**
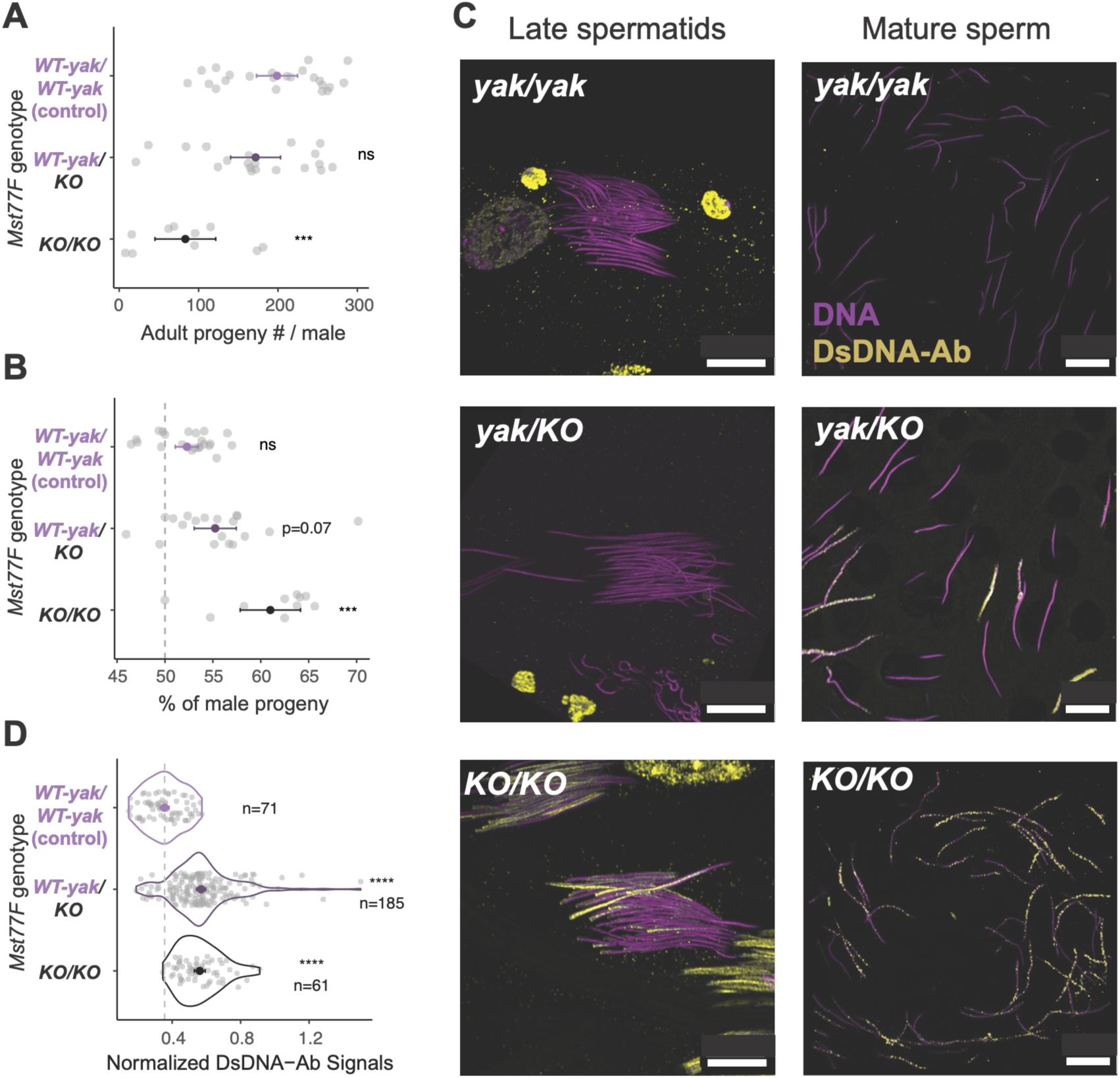
*Mst77F* knockouts in *D. yakuba* are fertile but have sex-ratio distortion. **(A)** Fertility of *D. yakuba* males carrying no, one, or two copies of wild-type *Mst77F-yak*. **(B)** Adult progeny sex ratios from *D. yakuba* males carrying no, one, or two copies of *Mst77F-yak.* Deviation from a 50:50 Mendelian expectation was measured using a χ² test. **(C)** Late spermatids and mature sperm from different *D. yakuba* males were stained with an α-dsDNA antibody (yellow) to assess their degree of chromatin compaction. Scale bar = 10 μm for all images. **(D)** Quantification of α-dsDNA antibody staining in sperm from wild-type, hemizygous, and *Mst77F-yak* knockout *D. yakuba* males. Statistical analyses were carried out using unpaired Student’s *t*-tests. (*p<0.05; ***p<0.001; ****p<0.0001; ns, not significant).

We found that *D. yakuba* males lacking one or both copies of *Mst77F-yak* produced male-biased offspring in a dose-dependent manner: homozygous knockout males show a more pronounced bias than hemizygous males (Fig. 6B). Consistent with our findings in *D. melanogaster*, we found that the depletion or absence of *Mst77F-yak* in *D. yakuba* males led to sperm DNA compaction defects (Fig. 6C–D) and abnormal nuclear size in mature sperm (Fig. S7B). While these compaction defects were pervasive, a subset of mature sperm remained viable and competent for fertilization, mirroring the incomplete penetrance of condensation failure observed in a particular *SD* background (*47*).

These phenotypes of *D. yakuba*, including the male-biased offspring, are reminiscent of those observed in crosses involving either hemizygous *Mst77F-mel* or homozygous *Mst77F-yak D. melanogaster* males, both of which are fertile yet exhibit sex-ratio distortion (Fig. 2). These findings suggest that both endogenous copies of *Mst77F-yak* are required to suppress delays or defects in sperm DNA compaction that would otherwise distort the progeny sex ratio in *D. yakuba*. Given the resulting male-biased progeny, we infer that these chromatin condensation defects also preferentially affect X-chromosome-bearing spermatids and mature sperm, as in *D. melanogaster*. Based on our detailed investigation of *Mst77F* in both *D. melanogaster* and *D. yakuba* males, we conclude that its rapid evolution across *Drosophila* species is more likely to be driven by its newly discovered role in suppressing meiotic drive than by its previously described essential function in male fertility.

## Discussion

Novel biological pressures shaping cellular processes can lead to unexpected signatures of genetic innovation, even in genes critical for viability or fertility (*48*). Despite their essential roles in male fertility in mammals and *Drosophila* species (*2*), protamine genes are among the most rapidly evolving protein-coding genes in many animals. Yet, the underlying causes of protamine evolution have remained unclear. Here, using genetic and cytological tools available in *D. melanogaster* and related species, we performed *in vivo* gene replacements to show that the rapid evolution of *Mst77F* protamine, which is essential for male fertility in *D. melanogaster*, ensures proper DNA condensation in sperm carrying either X or Y chromosomes during spermatogenesis. Failure to do so results in decreased fertility and sex-ratio distortion. Our findings represent the first experimental investigation into the causes underlying the rapid evolution of protamines in any species. They indicate that suppression of competition between X and Y chromosomes may be a primary driver of the rapid evolution of protamines.

Our findings indicate that the *Mst77F* protamine, which plays a crucial role in sperm genome compaction, may suppress meiotic drive by ensuring balanced representation of X and Y chromosomes in mature sperm. Disruptions in *Mst77F* dosage or sequence predominantly affect X chromosomes carrying specific targets, enabling Y-bearing sperm to outcompete X-bearing sperm and skew sex ratios. In contrast, a complete loss of *Mst77F* impairs condensation of all chromosomes, leading to complete male sterility in *D. melanogaster*. *Mst77F* might compact and directly protect an X-linked target from the drive. Y-linked duplications and expansions of protamines, including *Mst77Y* in *D. melanogaster* (*2*) or *Mst33A* in *D. yakuba* (*2*), may interfere with *Mst77F* function, favoring their transmission by disrupting X-bearing sperm and prompting rapid evolutionary adaptation of *Mst77F* to evade such antagonism and maintain sex-ratio parity. Alternatively, *Mst77F* might suppress sex-chromosome-linked sperm killers, which are selfish elements that bias their own transmission by disabling sperm carrying the opposing sex chromosome.

Such drivers need not reside on the Y. Indeed, our previous phylogenomic survey suggested that protamine-derived killers can arise on either the X or the Y, with the homologous sex chromosome serving as the target in each case (*2*). In this view, an autosomal protamine such as *Mst77F* is well positioned to act as a neutral arbiter, suppressing whichever sex chromosome gains a transmission advantage and thereby preserving the Fisherian 50:50 sex ratio that benefits the rest of the genome (*30, 31*). Loss of *Mst77F* could unleash these selfish elements, distort sex ratios, and cause sterility when genome-wide condensation fails. Thus, the widespread and rapid evolution of *Mst77F* and other protamines might reflect an ongoing evolutionary arms race with such sex-linked drivers. Future studies investigating the relationship between protamine evolution and the presence of sperm killers may illuminate the mechanisms underlying selfish genetic elements and their effects on male reproductive success and sperm chromatin integrity.

Our results, together with the genomic distribution of protamine genes, suggest that distinct protamines may have evolved to counter distinct meiotic drivers. *Mst77F* suppresses a sex-chromosome drive system whose target is X-linked, whereas the autosomal protamines *ProtA* and *ProtB* are implicated in suppressing the autosomal driver *Segregation Distorter* (*SD*) (*38*), but do not affect sex chromosome meiotic drive. Because the transmission conflict that fuels drive is most acute between the sex chromosomes, we expect sex-chromosome drive, which targets either the X or the Y in different lineages, to be a particularly recurrent pressure shaping protamine evolution (*2*).

Our findings reveal that both *Mst77F-mel* and *Mst77F-yak* suppress sex-chromosome meiotic drive in their respective species. In contrast to their roles in male fertility, both genes are haploinsufficient to suppress sex-ratio distortion, even in the presence of other protamines.

Moreover, loss of competition between X- and Y-bearing sperm due to a rare evolutionary X^Y fusion in the *montium* group of *Drosophila* species correlates with the loss of *Mst77F* and other protamines (*2*). These findings suggest that the retention, dosage, and rapid evolution of *Mst77F* and likely other protamine genes are associated with their role in suppressing meiotic drive during spermatogenesis. Our work highlights the power of an *in vivo* gene replacement strategy to investigate long-standing conundrums about the biological causes and consequences of the rapid evolution of genes essential for fertility and viability. It also highlights that, despite its ubiquity, Mendelian inheritance is ultimately a highly fragile truce.

## Acknowledgments

We thank Ritvija Agrawal, Sue Hammoud, Grant King, Amanda Larracuente, Pravrutha Raman, Maria Toro Moreno, and Janet Young for their valuable comments and suggestions that helped improve the manuscript. We are grateful to Dr. Barbara Wakimoto and members of the Malik, Ahmad, and Henikoff labs for fruitful discussions throughout the project. We also thank Benjamin Loppin, David Stern, and the Bloomington Drosophila Stock Center (supported by NIH P40OD018537) for the *Drosophila* strains, Lisa Kursel for the cDNA samples from *D. eugracilis* and *D. takahashii,* and Flybase (supported by NHGRI Award U41HG000739) for helping build bioinformatic tools across various *Drosophila* species’ genomes. This article is subject to HHMI’s Open Access to Publications policy. HHMI lab heads have previously granted a nonexclusive CC BY 4.0 license to the public and a sublicensable license to HHMI in their research articles. Pursuant to those licenses, the author-accepted manuscript of this article can be made freely available under a CC BY 4.0 license immediately upon publication.

## Funding

Damon-Runyon Cancer Research Foundation postdoctoral fellowship DRG 2438-21 (C-HC)

Taiwan National Science and Technology Council (NSTC 115-2311-B-001 -002 -MY3 to C-H C)

Academia Sinica intramural fund (C-HC)

National Institutes of Health grant R01-GM74108 (HSM) Howard Hughes Medical Institute Investigator (HSM)

## Author contributions

Conceptualization: CHC, HSM

Methodology: CHC, AC, INM, AN, HSM

Investigation: CHC, AC, HYL, INM, AN, EL, NL, HSM

Visualization: CHC, AC, INM, AN, HSM

Funding acquisition: CHC, HSM

Supervision: CHC, HSM

Writing – original draft: CHC, HSM

Writing – review & editing: CHC, AC, INM, AN, HSM

## Data and materials availability

Raw sequencing data were deposited in the NCBI Sequence Read Archive under BioProject accession no. PRJNA1290820. All data are available in the main text or the supplementary materials.

## Supplementary Materials

Materials and Methods

Figs. S1 to S8

Data S1 to S5

## References and Notes

1. P. B. Talbert, S. Henikoff, Histone variants at a glance. J Cell Sci 134, (2021).

2. C. H. Chang, I. Mejia Natividad, H. S. Malik, Expansion and loss of sperm nuclear basic protein genes in Drosophila correspond with genetic conflicts between sex chromosomes. Elife 12, (2023).

3. R. Balhorn, The protamine family of sperm nuclear proteins. Genome Biol 8, 227 (2007).

4. A. Torok, S. G. Gornik, Sperm Nuclear Basic Proteins of Marine Invertebrates. Results Probl Cell Differ 65, 15–32 (2018).

5. J. M. Eirin-Lopez, J. Ausio, Origin and evolution of chromosomal sperm proteins. Bioessays 31, 1062–1070 (2009).

6. N. Saperas et al., A unique vertebrate histone H1-related protamine-like protein results in an unusual sperm chromatin organization. FEBS J 273, 4548–4561 (2006).

7. N. Saperas, J. Ausio, Sperm nuclear basic proteins of tunicates and the origin of protamines. Biol Bull 224, 127–136 (2013).

8. S. M. Gartner et al., The HMG-box-containing proteins tHMG-1 and tHMG-2 interact during the histone-to-protamine transition in Drosophila spermatogenesis. Eur J Cell Biol 94, 46–59 (2015).

9. S. Chai et al., Coevolution and Adaptation of Transition Nuclear Proteins and Protamines in Naturally Ascrotal Mammals Support the Black Queen Hypothesis. Genome Biol Evol 16, (2024).

10. L. Luke, P. Campbell, M. Varea Sanchez, M. W. Nachman, E. R. Roldan, Sexual selection on protamine and transition nuclear protein expression in mouse species. Proc Biol Sci 281, 20133359 (2014).

11. G. J. Wyckoff, W. Wang, C. I. Wu, Rapid evolution of male reproductive genes in the descent of man. Nature 403, 304–309 (2000).

12. E. E. K. Kopania et al., Sperm competition intensity shapes divergence in both sperm morphology and reproductive genes across murine rodents. Evolution 79, 11–27 (2024).

13. S. Lupold et al., How sexual selection can drive the evolution of costly sperm ornamentation. Nature 533, 535–538 (2016).

14. C. Cho et al., Protamine 2 deficiency leads to sperm DNA damage and embryo death in mice. Biol Reprod 69, 211–217 (2003).

15. C. Cho et al., Haploinsufficiency of protamine-1 or -2 causes infertility in mice. Nat Genet 28, 82–86 (2001).

16. L. Moritz et al., Sperm chromatin structure and reproductive fitness are altered by substitution of a single amino acid in mouse protamine 1. Nat Struct Mol Biol 30, 1077–1091 (2023).

17. A. G. Clark, A. Civetta, Evolutionary biology. Protamine wars. Nature 403, 261, 263 (2000).

18. L. Luke, M. Tourmente, H. Dopazo, F. Serra, E. R. Roldan, Selective constraints on protamine 2 in primates and rodents. BMC Evol Biol 16, 21 (2016).

19. L. Luke, M. Tourmente, E. R. Roldan, Sexual Selection of Protamine 1 in Mammals. Mol Biol Evol 33, 174–184 (2016).

20. A. P. Rooney, J. Zhang, M. Nei, An unusual form of purifying selection in a sperm protein. Mol Biol Evol 17, 278–283 (2000).

21. S. Jayaramaiah Raja, R. Renkawitz-Pohl, Replacement by Drosophila melanogaster protamines and Mst77F of histones during chromatin condensation in late spermatids and role of sesame in the removal of these proteins from the male pronucleus. Mol Cell Biol 25, 6165–6177 (2005).

22. Z. Eren-Ghiani, C. Rathke, I. Theofel, R. Renkawitz-Pohl, Prtl99C Acts Together with Protamines and Safeguards Male Fertility in Drosophila. Cell Rep 13, 2327–2335 (2015).

23. T. Yamaki, G. K. Yasuda, B. T. Wakimoto, The Deadbeat Paternal Effect of Uncapped Sperm Telomeres on Cell Cycle Progression and Chromosome Behavior in Drosophila melanogaster. Genetics 203, 799–816 (2016).

24. R. Dubruille et al., Histone removal in sperm protects paternal chromosomes from premature division at fertilization. Science 382, 725–731 (2023).

25. J. Vedanayagam, C. J. Lin, E. C. Lai, Rapid evolutionary dynamics of an expanding family of meiotic drive factors and their hpRNA suppressors. Nat Ecol Evol 5, 1613–1623 (2021).

26. C. A. Muirhead, D. C. Presgraves, Satellite DNA-mediated diversification of a sex-ratio meiotic drive gene family in Drosophila. Nat Ecol Evol 5, 1604–1612 (2021).

27. F. J. Krsticevic, C. G. Schrago, A. B. Carvalho, Long-Read Single Molecule Sequencing to Resolve Tandem Gene Copies: The Mst77Y Region on the Drosophila melanogaster Y Chromosome. G3 (Bethesda) 5, 1145–1150 (2015).

28. J. Ricchio, F. Uno, A. B. Carvalho, New Genes in the Drosophila Y Chromosome: Lessons from D. willistoni. Genes (Basel) 12, (2021).

29. J. I. Park, G. W. Bell, Y. M. Yamashita, Derepression of Y-linked multicopy protamine-like genes interferes with sperm nuclear compaction in D. melanogaster. Proc Natl Acad Sci U S A 120, e2220576120 (2023).

30. R. A. Fisher, Chapter 6: Sexual Reproduction and Sexual Selection § Natural Selection and the sex-ratio. The Genetical Theory of Natural Selection (Clarendon Press, Oxford, UK, 1930), pp. 141.

31. W. D. Hamilton, Extraordinary sex ratios. A sex-ratio theory for sex linkage and inbreeding has new implications in cytogenetics and entomology. Science 156, 477–488 (1967).

32. X. Zhang et al., Broad phosphorylation mediated by testis-specific serine/threonine kinases contributes to spermiogenesis and male fertility. Nat Commun 14, 2629 (2023).

33. C. Rathke et al., Distinct functions of Mst77F and protamines in nuclear shaping and chromatin condensation during Drosophila spermiogenesis. Eur J Cell Biol 89, 326–338 (2010).

34. S. Kimura, B. Loppin, The Drosophila chromosomal protein Mst77F is processed to generate an essential component of mature sperm chromatin. Open Biol 6, (2016).

35. S. Kumar, G. Stecher, M. Suleski, S. B. Hedges, TimeTree: A Resource for Timelines, Timetrees, and Divergence Times. Mol Biol Evol 34, 1812–1819 (2017).

36. N. Raychaudhuri et al., Transgenerational propagation and quantitative maintenance of paternal centromeres depends on Cid/Cenp-A presence in Drosophila sperm. PLoS Biol 10, e1001434 (2012).

37. C. M. Schroeder et al., An actin-related protein that is most highly expressed in Drosophila testes is critical for embryonic development. Elife 10, (2021).

38. L. F. Gingell, J. R. McLean, A Protamine Knockdown Mimics the Function of Sd in Drosophila melanogaster. G3 (Bethesda) 10, 2111-2115 (2020).

39. M. Herbette et al., Distinct spermiogenic phenotypes underlie sperm elimination in the Segregation Distorter meiotic drive system. PLoS Genet 17, e1009662 (2021).

40. N. L. Nuckolls et al., wtf genes are prolific dual poison-antidote meiotic drivers. Elife 6, (2017).

41. L. Caro et al., An animal toxin-antidote system kills cells by creating a novel cation channel. PLoS Biol 23, e3003182 (2025).

42. E. Ben-David, A. Burga, L. Kruglyak, A maternal-effect selfish genetic element in Caenorhabditis elegans. Science 356, 1051–1055 (2017).

43. Y. Hua et al., Structural duality enables a single protein to act as a toxin-antidote pair for meiotic drive. Proc Natl Acad Sci U S A 121, e2408618121 (2024).

44. S. You et al., A toxin-antidote system contributes to interspecific reproductive isolation in rice. Nat Commun 14, 7528 (2023).

45. D. L. Brutlag, Molecular arrangement and evolution of heterochromatic DNA. Annu Rev Genet 14, 121–144 (1980).

46. C. Drosophila 12 Genomes et al., Evolution of genes and genomes on the Drosophila phylogeny. Nature 450, 203–218 (2007).

47. J. T. Ridges et al., Selfish chromosomes exploit a germline checkpoint to eliminate competing gametes. Nat Commun 17, 1532 (2026).

48. C. L. Brand, M. T. Levine, Functional Diversification of Chromatin on Rapid Evolutionary Timescales. Annu Rev Genet 55, 401–425 (2021).

49. N. Kost et al., Multimerization of Drosophila sperm protein Mst77F causes a unique condensed chromatin structure. Nucleic Acids Res 43, 3033–3045 (2015).

