## Supplementary material for "Rapid protamine evolution suppresses meiotic drive in *Drosophila*": Sup Materials

### Methods & Materials

<u>CRISPR/Cas9 knockout of *Mst77F* in *D. melanogaster* and *D. yakuba*</u>

To generate the *Mst77F* knockout strain, we adapted the *pCFD4-U6:1_U6:3 tandem gRNAs* plasmid (Addgene #49411) with two guide RNAs (gRNAs) targeting either the 5' or 3' end of *Mst77F*. For the repair construct, we amplified four fragments via PCR: the 3×P3 *DsRed* marker, the backbone of pDsRed-attP (Addgene #51019), and ~1 kb of homologous sequences flanking the upstream and downstream regions of *Mst77F*. These four fragments were then annealed using NEBuilder® HiFi DNA Assembly Master Mix into one 'repair construct.' We used the Q5® Site-Directed Mutagenesis Kit (NEB catalog E0554S) to mutate the gRNA target PAM sites on the repair construct to avoid recutting of the repair construct by CRISPR/Cas9 and the gRNAs.

For *D. melanogaster*, these two constructs (one encoding the repair construct and the other encoding the guide RNAs) were co-injected into the *y[1] M{GFP[E.3×P3]=vas-Cas9.DsRed-}ZH-2A w[1118]* strain (BDSC stock #55821) embryos by BestGene Inc. The transgenic flies were backcrossed to the *yw* strain, selected using the DsRed marker, and then balanced using the *TM3, Sb* balancer chromosome. We used PCR to validate the successful *Mst77F-mel* knockout (Fig. S1B). The *Mst77F-mel* knockout was further confirmed using complementation tests with the *Mst77FΔ1* knockout flies (*34*) (kind gift from Dr. Benjamin Loppin)(Fig. S1B).

For *D. yakuba,* we similarly constructed gRNA and repair template plasmids as described above. These two constructs were co-injected into a *D. yakuba* strain encoding *nanos-cas9* on the X chromosome (*41*) (kind gift from Dr. David Stern) by Genetivision Inc. The transgenic flies were backcrossed to a *w* strain of *D. yakuba* and selected using the *DsRed* marker (Fig. S9A). We used PCR to validate the successful *Mst77F-yak* knockout (Fig. S9B). The *D. yakuba* stocks are maintained as heterozygotes by selecting the *DsRed* marker every 2–3 generations. All primers are listed in Data S1.

<u>Recombinase-mediated *in vivo* gene replacements at the endogenous *Mst77F* locus in *D. melanogaster*</u>

For the gene replacement and rescue experiments in *D. melanogaster*, we utilized the attP site in the integrated repair construct of *Mst77F KO* flies. First, we removed the *DsRed* marker in *Mst77F KO* flies by crossing the knockout strain with *Cre* recombinase-carrying males (BDSC #1501). This removes the *DsRed* gene, which is flanked by *loxP* sites. The resulting *Mst77F KO/TM6B, Cre* males were backcrossed to the maternal *Mst77F KO/TM3, Sb* strain. The resulting *DsRed*-negative flies carrying the *TM3, Sb* balancer were selected to establish a stable stock.

Next, we amplified the coding sequences of different *Mst77F* orthologs from various species using reverse transcription PCR (RT-PCR) from male RNA. We cloned these *Mst77F* copies flanked by the promoter and untranslated regions from *D. melanogaster*, into a vector carrying attB and 3×P3 DsRed (kind gift from Dr. Courtney Schroeder) using NEBuilder® HiFi DNA Assembly Master Mix.

To generate the overexpression construct, the PiggyBAC backbone (kind gift from Dr. Courtney Schroeder), containing GFP and mini-white markers, was linearized via double digestion with SbfI-HF and NotI-HF. Subsequently, the Mst77F-yak transgene flanked by *D. melanogaster* promoter and UTR sequences was subcloned from the attB donor vector into the linearized PiggyBAC vector using NEBuilder® HiFi DNA Assembly Master Mix.

These constructs were co-injected along with a helper plasmid encoding either ΦC31 integrase or *piggyBac* transposase into *Mst77F-mel KO [DsRed-flox]/TM3, Sb* flies by BestGene Inc. The resulting transgenic flies were backcrossed with *Mst77F-mel KO [RFP-flox]/TM3, Sb* to ensure an identical genetic background. The resulting *DsRed*-positive flies were selected and maintained using the *TM3, Sb* balancer chromosome. All primers are listed in Data S1.

<u>Constructing chimeras between *Mst77F-mel and Mst77F-yak*</u>

We used PCR to amplify different regions of *Mst77F-mel* and *Mst77F-yak* using the plasmids created in the previous section. We then constructed attB clones with the *Mst77F-mel/Mst77F-yak* chimeras by annealing the fragments using NEBuilder® HiFi DNA Assembly Master. In this way, we generated vectors carrying attB, 3×P3 DsRed, and different *Mst77F-mel/Mst77F-yak* chimeras. These constructs were again injected into *Mst77F-mel KO [DsRed-flox]/TM3, Sb* flies by BestGene. The resulting transgenic flies were backcrossed with *Mst77F-mel KO [DsRed-*

*flox]/TM3, Sb* to ensure an identical genetic background. *DsRed*-positive flies were selected and maintained using TM3, Sb balancer chromosomes. All primers are listed in Data S1.

Sequencing and RT-PCR

We extracted genomic DNA from individual flies by grinding a single fly in a buffer containing 10 mM Tris-HCl (pH 8), 1 mM EDTA, 25 mM NaCl, and 200 μg/mL Proteinase K. The mixture was incubated at 37°C for 30 minutes, followed by heat inactivation of Proteinase K at 95°C for 3 minutes. For RNA extraction, we homogenized flies or specific tissues in TRIzol reagent (Invitrogen), added chloroform, and followed the manufacturer's protocol for RNA extraction using the RNA Clean & Concentrator kit (Zymo Research) with DNase I to purify the RNA. Samples were treated with DNase I (Zymo Research) and further purified and concentrated using the same kit if there was still DNA contamination. The resulting RNA was reverse-transcribed into cDNA using SuperScript III First-Strand Synthesis (Invitrogen Inc.). All primers are listed in Data S1.

Cross schemes and Fertility assays

To visualize histone removal, we introduced *His2AV-mRFP* and *cid-GFP* from BDSC strain #91708 into *Mst77F*-replacement backgrounds. Females from the reporter strain were crossed to males carrying either the *Mst77F-mel* or *Mst77F-yak* transgene. Resulting virgin females (*His2AV-mRFP, cid-GFP / Mst77F*) were backcrossed to the paternal transgenic lines. Finally, progeny carrying both fluorescent reporters and the transgenic *Mst77F* allele balanced over *TM3, Sb* were selected to establish stable stocks. A similar cross was used to increase the copy number of *Mst77F-yak* on the third chromosome.

To introduce attached X^Y chromosomes into *Mst77F*-deficient backgrounds, we utilized a strain carrying both attached X^X and attached X^Y chromosomes (BDSC strain #9460). We first crossed males from our *Mst77F-KO* or *Mst77F-yak* strain to females from #9460. The resulting X^X/Y females were then backcrossed to #9460 males to get X^Y/Y males carrying the *Mst77F-KO* or *Mst77F-yak*, identified by their respective DsRed selection markers. These X^Y/Y males were backcrossed once more to #9460 females again to segregate out the extra Y chromosome, yielding X^Y males carrying the desired *Mst77F* allele. We maintain these experimental stocks by selecting for the DsRed marker in each generation.

To test whether the *Zhr* locus (comprising the 359-bp satellite repeats) is the target locus of *Mst77F*-regulated meiotic drive, we genetically removed the *Zhr* region from the *Mst77F* transgenic background. First, the X chromosome from the strain (BDSC strain #25140) with a deletion of *Zhr* (Δ*Zhr*) was introduced to the TM3/TM6B double-balancer background to generate a stable base stock, Δ*Zhr;* TM3/TM6B. To introduce the specific Mst77F variants, Δ*Zhr;* TM3/TM6 females were crossed to males from the *Mst77F-KO/*TM3, *Mst77F-mel/*TM3 or *Mst77F-yak/*TM3 stock. The resulting Δ*Zhr; Mst77F-KO/*TM3, Δ*Zhr;Mst77F-mel/*TM3 or Δ*Zhr; Mst77F-yak/*TM3 males were backcrossed to the baseline to establish balanced stocks (Δ*Zhr; Mst77F-KO/*TM3, Δ*Zhr; Mst77F-mel/*TM3, and Δ*Zhr; Mst77F-yak/*TM3). To test potential dosage effect, we crossed the Δ*Zhr*; *Mst77F-mel* strain and the Δ*Zhr*; *Mst77F-KO* strain to yield trans-heterozygous experimental males (Δ*Zhr*; *Mst77F-mel/Mst77F-KO*). Finally, these males were crossed to *yw* or OreR females at a 1:5 ratio for 9 days in total, and their female and male progeny were scored.

To measure male fertility and progeny sex ratios, we crossed each 2–6-day-old tester male to five 2–6-day-old virgin females from the wild-type Oregon-R *D. melanogaster* strain, an attached X^X strain (BDSC strain #9460), or a *white D. yakuba* strain (kind gift from Dr. David Stern) at 25°C or 29°C. For *D. melanogaster,* we transferred the mating pairs to new vials every three days and counted all resulting offspring from the first nine days. For *D. yakuba,* we transferred the mating pairs to new vials every seven days and counted all resulting offspring from the first 21 days. All raw results are listed in Data S2.

Mst77F dephosphorylation assays and Western blot

To extract Mst77F, testes from 2-day-old virgin males were dissected in the chilled dissection buffer (1x Cutsmart buffer, 150 mM NaCl, 1x protease inhibitor). Samples were then lysed in the dissection buffer with 0.1% Triton X-100 through five freeze-thaw cycles using liquid $N_2$, followed by centrifugation at 15000xg for 10 minutes at 4 °C. The resulting supernatant was divided into equal aliquots and treated with either 5 units of Quick-CIP (NEB) or an equivalent volume of lysis buffer (control). Following a 1-hour incubation at 37°C with 700 rpm shaking, samples were mixed with 4x Laemmli buffer and heated at 90°C for 5 minutes.

Electrophoresis was performed using 12% SDS-PAGE at 100V, followed by a 1-hour transfer to nitrocellulose membranes at 100V. After blocking with Hyblock buffer for 5 minutes, the membrane was incubated overnight at 4°C with primary antibodies: mouse anti-FLAG (1:2000; Sigma F3165) and rabbit anti-α-tubulin (1:5000; Abcam ab52866) diluted in 5% BSA in TBST (0.05% Tween 20). Following three washes with TBST, the membrane was incubated with fluorescent secondary antibodies—Goat α-mouse (1:10,000; LI-COR 926-32210) and Goat α-rabbit (1:10,000; Abcam ab175773)—in 5% BSA in TBST for 1 hour at room temperature. After three final washes in TBST, the blots were visualized using an Invitrogen iBright FL1500 Imaging System. All raw results are listed in Data S3.

<u>Quantification of seminal vesicle sizes</u>

Testes with seminal vesicles were carefully dissected from 5-day-old virgin males in 1X PBS buffer and mounted in 30 µL of 1X PBS buffer to avoid squashing the tissue. Images were captured under bright-field illumination (ZEISS Scope A1), and the total 2D surface area (in $\mu m^2$) was manually outlined and calculated via ZEISS Zen 3.9. All raw results are listed in Data S4.

<u>Immunofluorescence assays</u>

For immunofluorescence, testes from 1–3-day-old virgin male flies were dissected and fixed in 4% paraformaldehyde (PFA) for 20 minutes at room temperature. The samples were permeabilized in 0.3% sodium deoxycholate in PBS with 0.1% Triton X-100 (PBST) for 15 minutes, repeated twice, followed by three washes with PBST. Testes were then blocked for 1 hour in 3% BSA in PBST (blocking buffer) and incubated overnight with primary antibodies (1:200 rabbit α-FLAG; Proteintech 0543-1-A, 1:1000 mouse α-DsDNA; Abcam 27156) diluted in the blocking buffer at

4°C. On the second day, the testes were washed three times with PBST and treated with corresponding secondary antibodies (1:1000) diluted in the blocking buffer at room temperature for 2 hours, with Hoechst 33342 (1:1000 Invitrogen) added for DNA staining. After three additional washes with PBST and a final wash with PBS, the testes were mounted onto slides using SLOWFADE antifade mounting medium with DAPI (Invitrogen Inc. S36938) for imaging.

Fluorescence *in situ* hybridization assays

For fluorescence in situ hybridization (FISH), testes were dissected from 1–3-day-old virgin male flies and fixed in 4% PFA for 30 minutes at room temperature. Similarly for immune-FISH, the samples from immunostaining were post-fixed in 4% PFA for 6 minutes at room temperature. The samples were treated with RNase A (Invitrogen) in PBST for 10 minutes. After treatment, the samples were washed twice with 2X saline sodium citrate with 0.1% Tween-20 (2X SSCT) and 1 mM EDTA, followed by a single wash with hybridization buffer. We then treated the samples with 50 µL of hybridization buffer containing 50–100 ng of probes (see Data S1) and 1 mM EDTA, and denatured the samples at 95 °C for 8 min before hybridizing them overnight at 37 °C in the cap of an Eppendorf tube sealed with heat-sealing film. The next day, after washing three times with 4X SSCT and 0.1X SSC, the testes were mounted onto slides using SLOWFADE antifade mounting medium with DAPI (Invitrogen Inc. S36938) for imaging. The solution and details of the FISH protocol followed the procedure outlined previously (*42*), except that we supplemented the 2X SSCT with 1 mM EDTA during hybridization (*43*).

Imaging analyses and sperm shape assays

We used a DragonFly spinning-disk confocal microscope with a 100X oil-immersion objective to capture all images, using Fusion software for acquisition. For the FISH experiment, sperm were categorized based on distinct probe signals. To measure sperm length and fluorescence intensity in immunostaining, we used the surface modeling function in IMARIS to identify long, straight sperm nuclei. Following an additional round of manual curation, we calculated sperm length by doubling the Ellipsoid Axis Length C value. To quantify signal intensity, we measured the fluorescence of the α-dsDNA antibody and normalized it to the Hoechst 33342 signal to correct for signal decay across the Z-axis. All raw results are listed in Data S5.

RNA sequencing and analyses

We collected 5–10 pairs of testes from 2–3-day-old virgin male flies and carefully removed the accessory glands on ice. Testes were homogenized in 200 μL TRIzol, followed by the addition of 40 μL chloroform. After centrifugation at 13,000 xg for 15 minutes at 4 °C, we collected 100 μL of the aqueous phase and purified the RNA using the Zymo RNA Clean & Concentrator kit (R1015) with on-column DNase I treatment. The RNA was dissolved in 20 μL $ddH_2O$. If Nanodrop or PCR detected DNA or phenol contamination, we repeated the purification using the same Zymo kit. Purified total RNA was submitted to Novogene for poly(A) enrichment, library preparation, and sequencing on the NovaSeq X platform, generating at least 6 Gb of 150-bp paired-end reads per sample. RNA sequence reads are in NCBI's SRA under BioProject PRJNA1290820.

To quantify the expression of *Mst77F* orthologs in *Mst77F* gene replacement males, we generated a custom transcriptome reference based on FlyBase r6.63, in which the original *Mst77F-mel* transcript sequence was removed and replaced with FLAG-tagged *Mst77F-mel* and *Mst77F-yak* sequences from the constructs. We then quantified transcript abundance using SALMON v1.10.1 (*44*) using the following parameters: --validateMappings --allowDovetail --recoverOrphans --gcBias --seqBias. For comparison, RNA-seq data from wild-type testes ($w^{1118}$ strain) were obtained from the publicly available dataset PRJNA687271 (*45*).

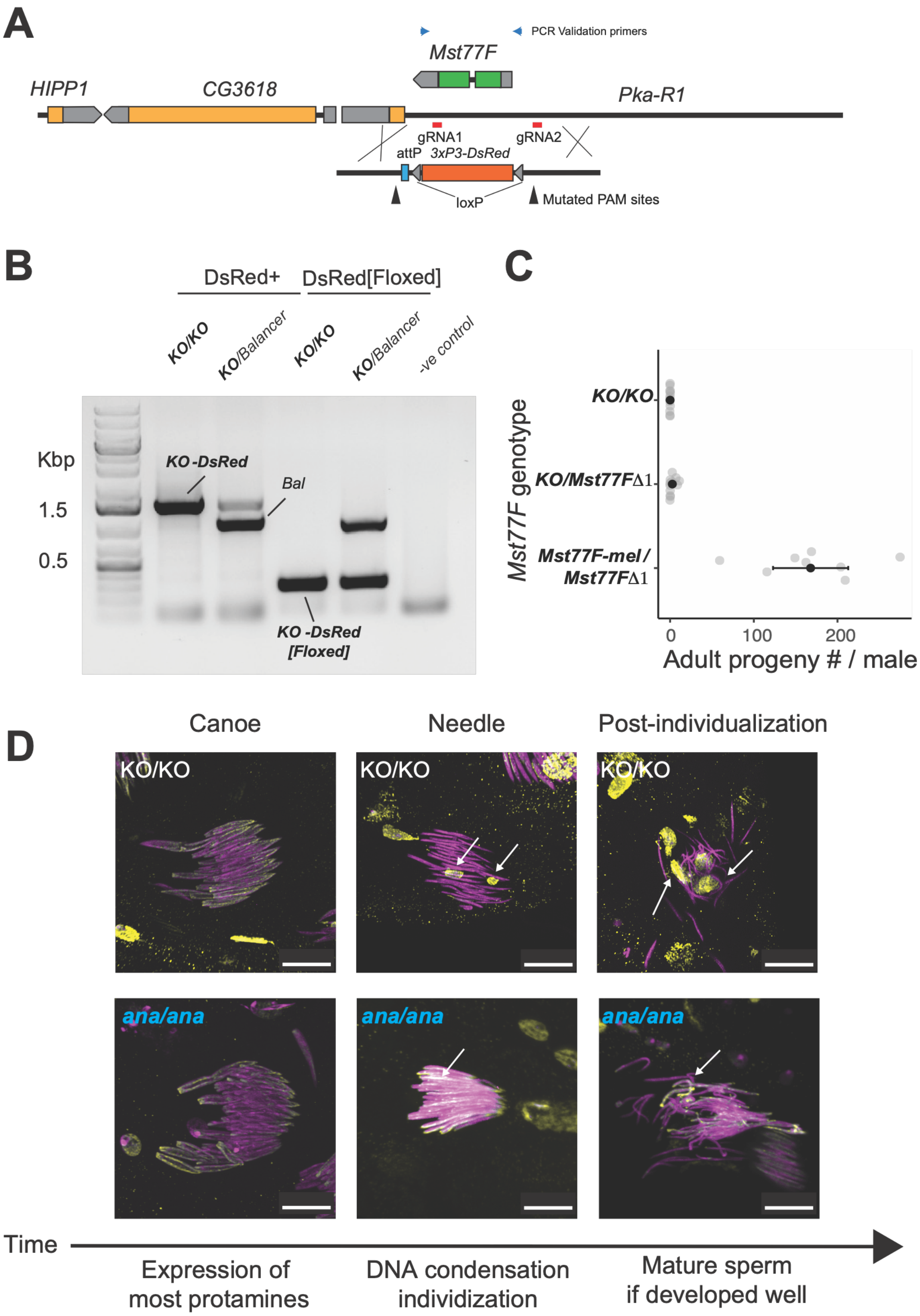


**Fig. S1: *Mst77F* knockout and replacement in *Drosophila melanogaster*. (A)** Schematic representation of the genomic region flanking the *Mst77F* locus in *D. melanogaster*. The
225 positions of the flanking *HIPP1*, *CG3618*, and *Pka-R1* genes are shown. The CRISPR/Cas9

strategy for generating the knockout (*KO*) allele is depicted, including the locations of guide RNAs gRNA1 and gRNA2, and a repair construct carrying a 3×P3-DsRed marker, an attP site, and loxP sites. PAM sites were mutated in the repair construct to prevent re-cutting by CRISPR/Cas9. PCR validation primer sites are shown as blue arrowheads. **(B)** PCR genotyping of flies carrying the *KO* and floxed alleles shows bands corresponding to the *Mst77F-mel KO*-DsRed, *KO*-DsRed/balancer (*Bal*), *KO*-DsRed[floxed], *KO*-DsRed[floxed]/balancer flies, as well as a negative control. **(C)** *KO/KO* males and trans-heterozygote *KO*/*Mst77FΔ1* (an independently derived *Mst77F* knockout allele(*34*)) are sterile. This male sterility can be complemented by FLAG-tagged *Mst77F-mel*, indicating that *Mst77F-mel* functions as a wild-type allele. **(D)** α-DsDNA antibody staining of sperm from the *KO*/*KO D. melanogaster* males. *KO*/*KO* cannot produce mature sperm and shows chromatin condensation defects in sperm nuclei (which leads to α-DsDNA antibody staining) during individualization and post-individualization stages of spermatogenesis. DNA was stained with Hoechst 33342 (magenta). Scale bar = 10 μm for all images.

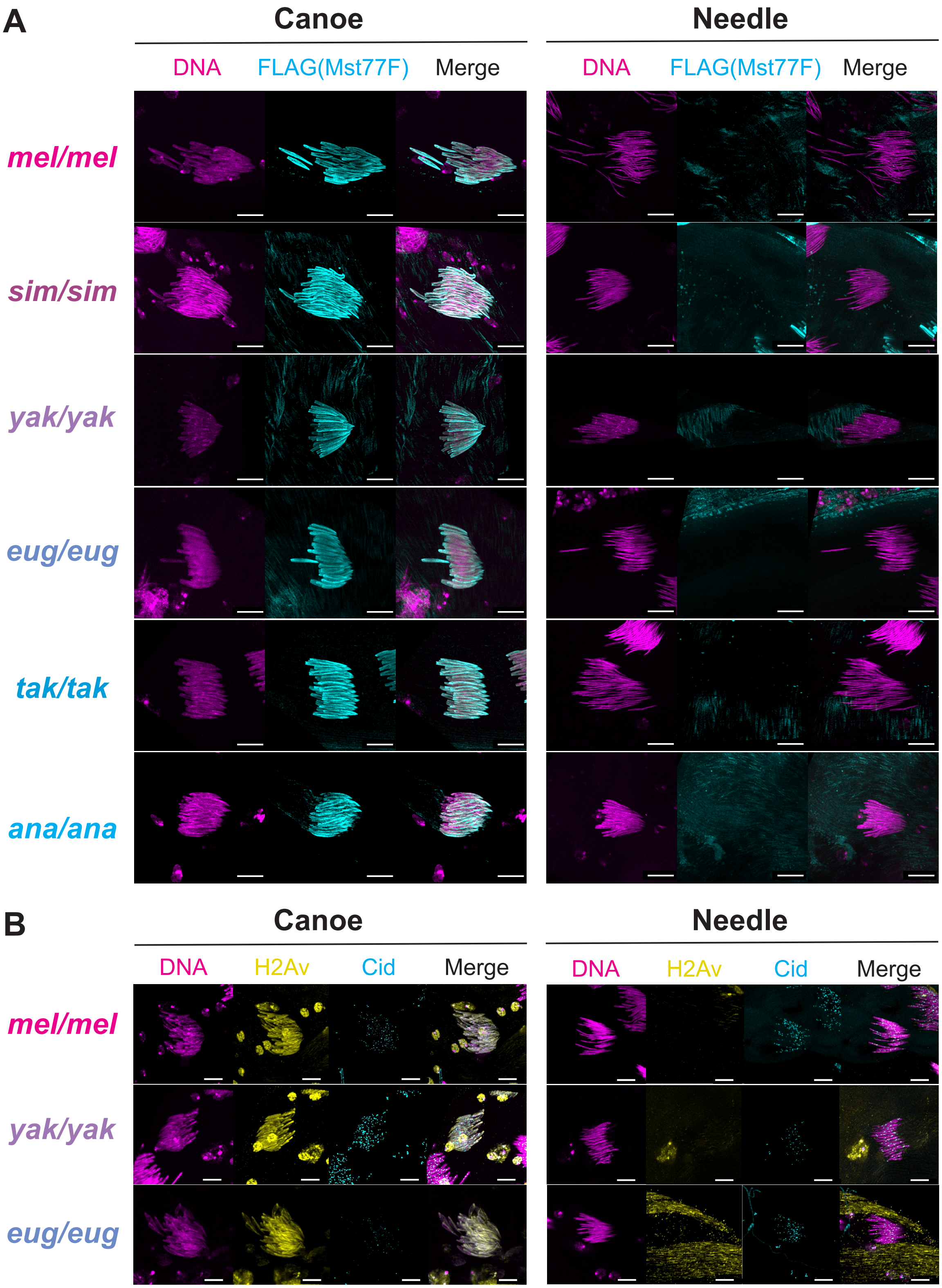
A
Canoe
Needle
DNA
FLAG(Mst77F)
Merge
DNA
FLAG(Mst77F)
Merge
mel/mel
sim/sim
yak/yak
eug/eug
tak/tak
ana/ana
B
Canoe
Needle
DNA
H2Av
Cid
Merge
DNA
H2Av
Cid
Merge
mel/mel
yak/yak
eug/eug

240

**Fig. S2: Expression and localization of *Mst77F* orthologs and histones during spermatid maturation stages.** Testes from transgenic flies expressing different *Mst77F* orthologs were immunostained and imaged at the Canoe and Needle (individualization) stages of spermatogenesis. **(A)** FLAG-tagged Mst77F was detected using α-FLAG antibody (cyan), DNA was stained with Hoechst 33342 (magenta). Mst77F is known to be expressed at the canoe stage and retained in mature sperm. However, during and after the needle stage, sperm nuclei are highly compacted, preventing antibody penetration. Signal detected outside spermatid nuclei reflects nonspecific binding. **(B)** Histone 2A variant (*His2Av*) and the centromeric histone (*cid*) were visualized via endogenously tagged *His2Av-RFP* (yellow) and *cid-GFP* (cyan). Scale bar = 10 μm for all images.

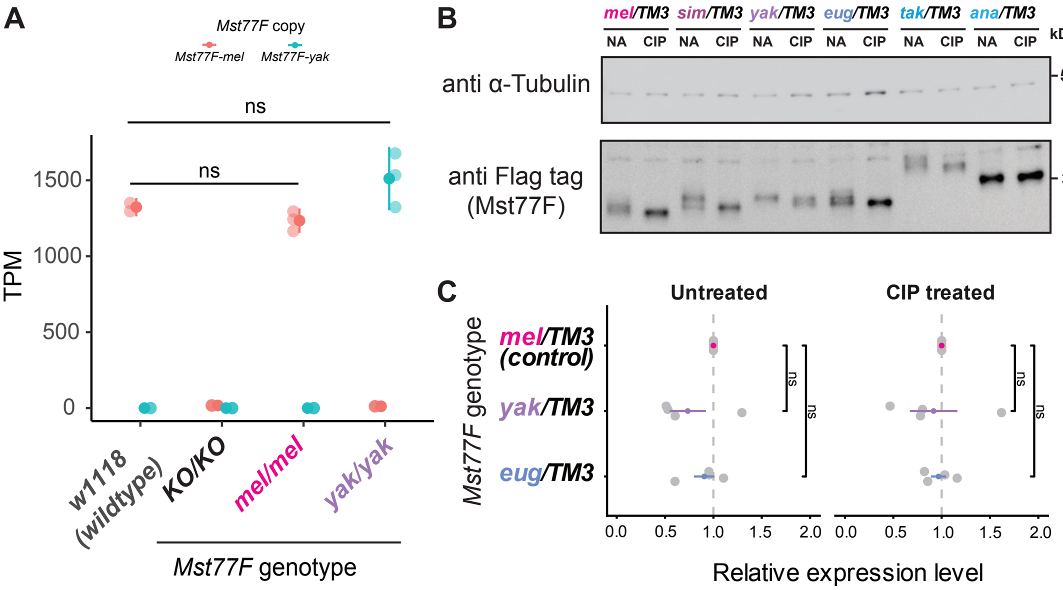


**Fig. S3. Expression levels and phosphorylation of *Mst77F* transgenes in testes.** (**A**) We quantified the expression of *Mst77F* in the testes of a wild-type strain (*w1118*), *an Mst77F* knockout strain (*KO/KO*), a rescued homozygous *Mst77F-mel* strain (*mel/mel*), and a homozygous *Mst77F-yak* replacement strain (*yak/yak*) using RNA-seq. To avoid mapping bias and distinguish the expression of *Mst77F-mel* and *Mst77F-yak*, we mapped the reads to a modified Flybase transcriptome reference, in which the original *Mst77F* transcript sequence was replaced with FLAG-tagged *Mst77F-mel* and *Mst77F-yak* sequences from the constructs we used. Statistical analyses were carried out using unpaired Student's *t*-tests, comparing with the wild-type flies. (**B**) Phosphorylation analysis via Western Blot of transgenic Mst77F-FLAG proteins across species. A downward shift in the anti-Flag signal following dephosphorylation treatment via Calf Intestinal Phosphatase (CIP) indicates phosphorylation of all Mst77F proteins except the most distant Mst77F-*ana*. α-tubulin serves as a loading control. (**C**) Quantification of three different Mst77F protein levels (normalized to α-tubulin) in the presence or absence of CIP treatment. Significance was determined by unpaired Student's *t*-tests (*p<0.05; **p<0.01; ***p<0.001; ****p < 0.0001; ns, not significant).

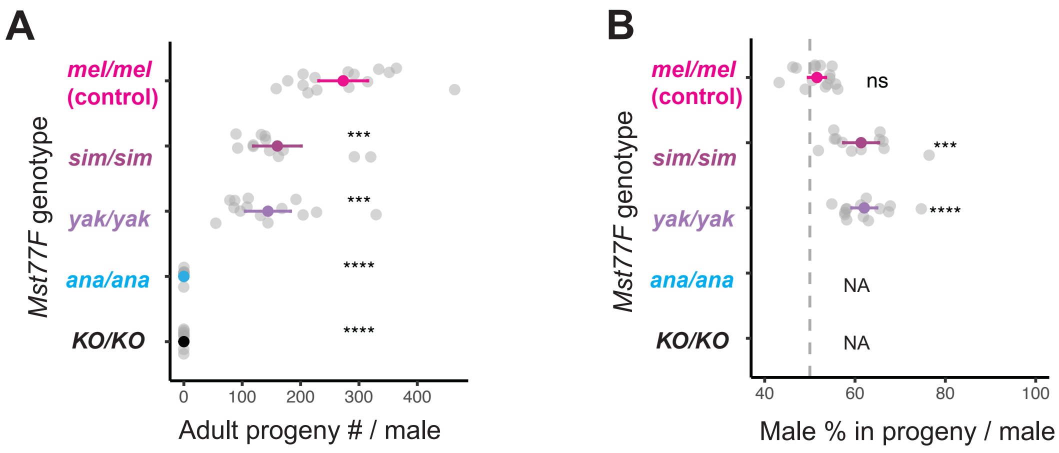


**Fig. S4: *Mst77F* gene replacements in *D. melanogaster* reduce male fertility and result in sex-ratio distortion at 29°C**. **(A)** Similar to 25°C, transgenic *D. melanogaster* males carrying two copies of *Mst77F-sim or Mst77F-yak* partially rescue fertility to levels comparable to *Mst77F-mel*. In contrast, flies carrying two copies of the *Mst77F-ana* are sterile, just like knockout (*KO*) flies. **(B)** Sex ratios of the adult progeny of transgenic *D. melanogaster* males carrying two copies of *Mst77F-mel, Mst77F-sim, or Mst77F-yak* mated to wild-type females. Each dot represents data from a single male. Transgenic *D. melanogaster* males carrying two copies of *Mst77F-mel* produce equal numbers of male and female offspring, consistent with Mendelian expectations. In contrast, *D. melanogaster* males carrying two copies of *Mst77F-sim* or *Mst77F-yak* produce significantly more male offspring. Each dot represents data from a single male. All statistics are carried out using unpaired Student's *t*-tests against the control or $\chi^2$ test against a 50:50 ratio (*p<0.05; **p<0.01; ***p<0.001; ****p < 0.0001; ns, not significant).

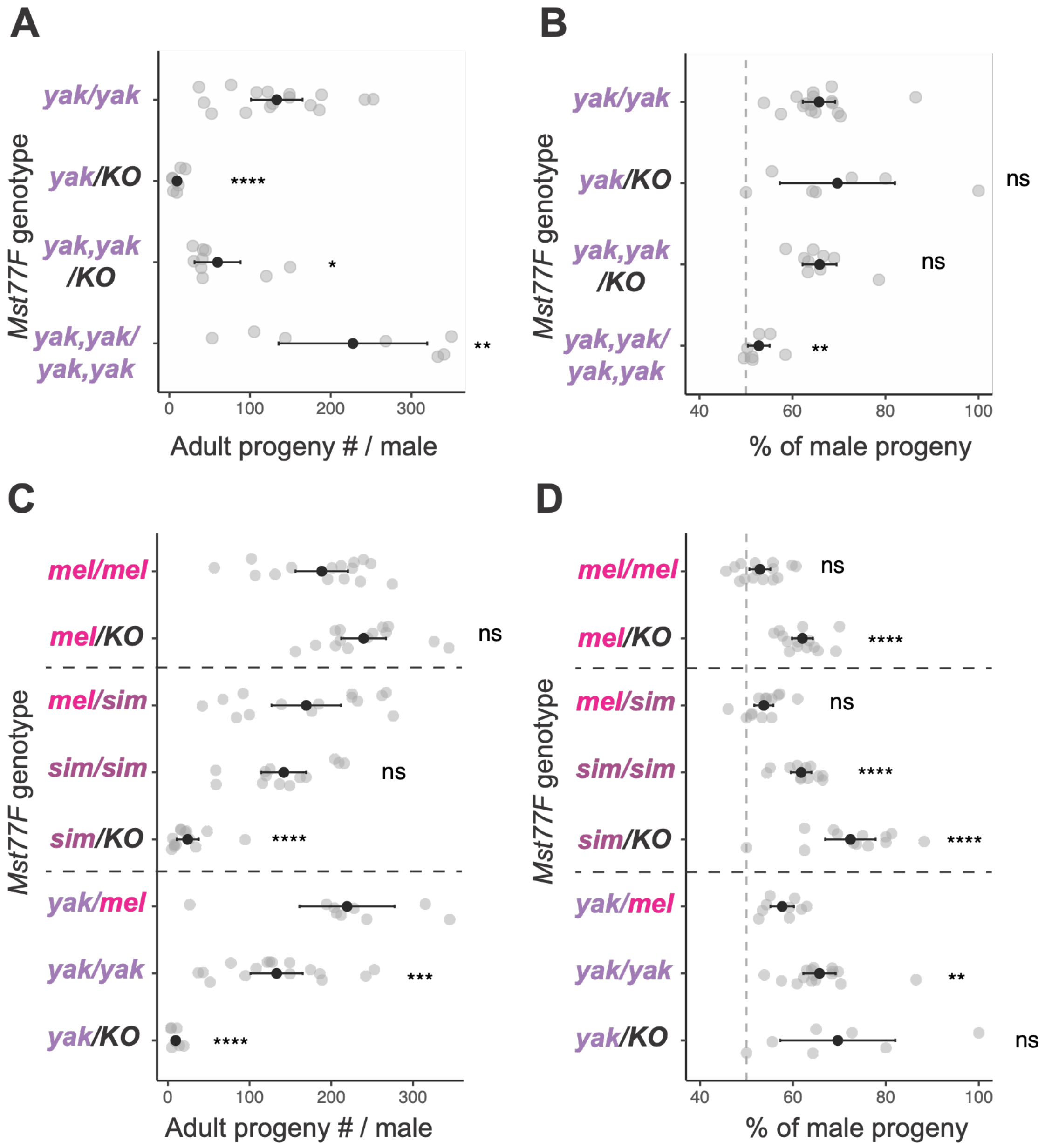


**Fig. S5: Increased dosage or endogenous *Mst77F-mel* expression rescues fertility defects and sex-ratio distortion driven by *Mst77F* ortholog replacements. (A)** Total adult progeny per male showing that supplementing with additional copies of *Mst77F-yak* rescues male fertility defects. **(B)** The percentage of male progeny from males carrying varying copy numbers of the *D. yakuba Mst77F* ortholog (*yak*) in an *Mst77F-mel* knockout (*KO*) background. An increased dosage of the *D. yakuba* transgene (*yak,yak/yak,yak*) significantly alleviates male-biased sex-ratio distortion toward the expected 50:50 baseline (indicated by the vertical dashed gray line). **(C)** Total adult progeny per male showing that supplementing with one additional copy of endogenous *Mst77F-mel* similarly restores male fertility. **(D)** The percentage of male progeny, demonstrating that the distorted male-biased sex ratios seen in hemizygous *Mst77F-sim* or *Mst77F-yak* males are alleviated by an additional copy of endogenous *Mst77F-mel*.

For all panels, horizontal dashed lines separate genetic groups and each dot represents data from a single male. Statistical comparisons for fertility **(A, C)** were evaluated using unpaired Student's *t*-tests, and sex ratios **(B, D)** were analyzed via $\chi^2$ test against 50:50 ratio (*$p<0.05$; **$p<0.01$; ****$p < 0.0001$; ns, not significant).

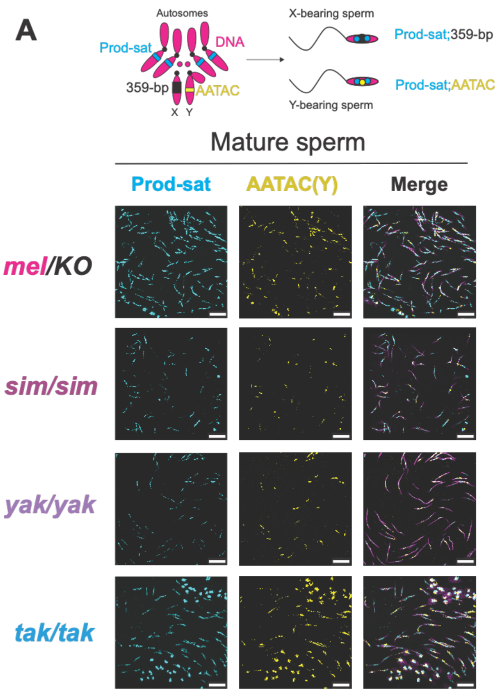


**Fig. S6: *Mst77F* gene replacements in *D. melanogaster* decrease the proportion of X-bearing sperm.** FISH was performed using the Y-linked AATAC (yellow) and Prod-sat (cyan) probes on seminal vesicles from homozygous *D. melanogaster* males carrying only one copy of *Mst77F-mel (mel/KO)* or two copies of *Mst77F* orthologs from other species (*sim/sim, yak/yak,* and *tak/tak)*.

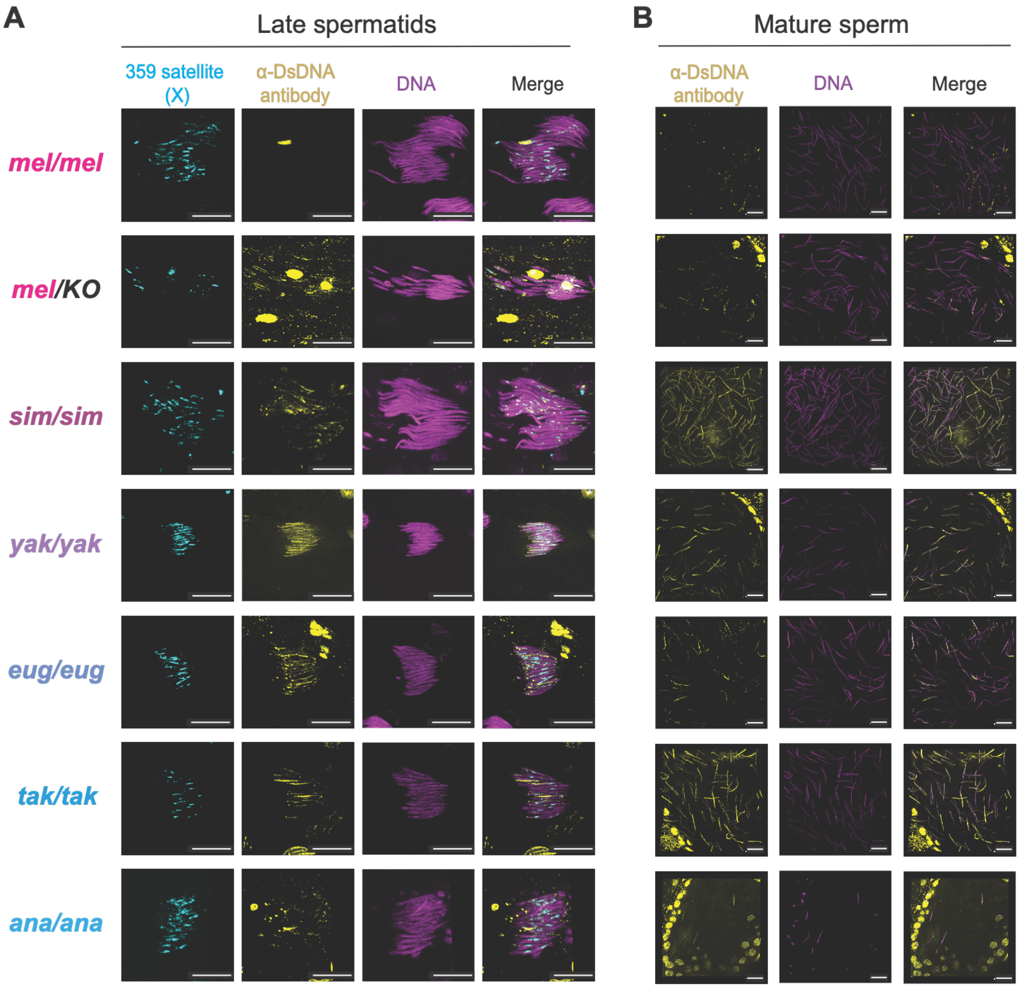


**Fig. S7: *Mst77F* sequence divergence and dosage affect sperm nuclei condensation of X-bearing sperm in *D. melanogaster*.** Late spermatids **(A)** and mature sperm **(B)** were stained with an α-dsDNA antibody (yellow) to assess their degree of chromatin compaction using immunostaining. DNA was stained with Hoechst 33342 (magenta). X-linked 359-bp satellite repeats were visualized with a DNA FISH probe (cyan). Late spermatids and mature sperm from flies carrying only one copy of *Mst77F-mel (mel/KO)* or two copies of *Mst77F* orthologs from other species (*sim/sim, yak/yak, eug/eug, tak/tak*, and *ana/ana)* exhibited chromatin compaction defects, as indicated by persistent α-dsDNA staining, whereas flies carrying two copies of *Mst77F-mel* (*mel/mel*) did not. In addition, flies with *Mst77F-ana/Mst77F-ana* have almost no mature sperm and are sterile. Scale bar = 10 µm for all images.

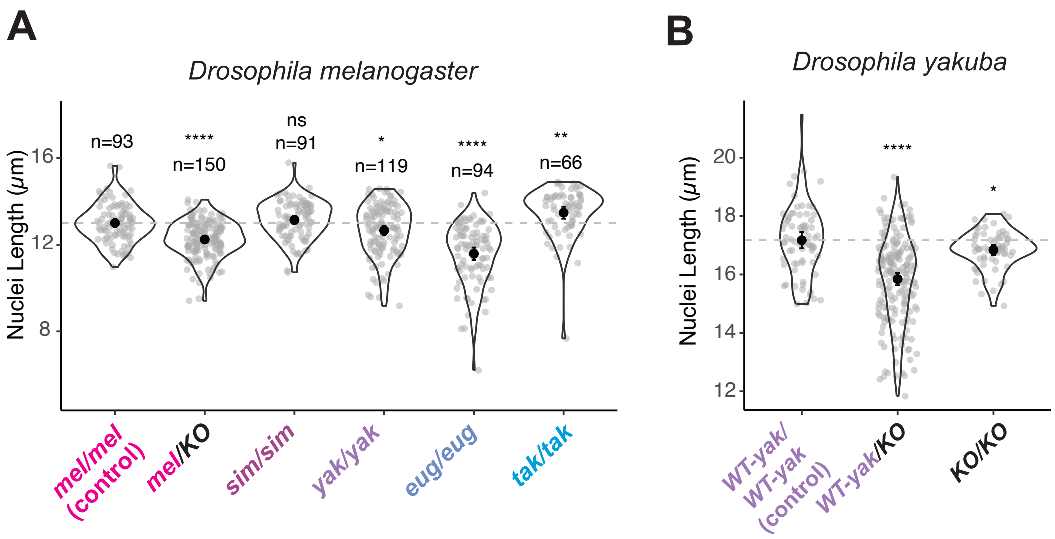


**Fig. S8: *Mst77F* sequence divergence and dosage affect sperm nuclei length in *D. melanogaster*.** (A) Sperm nuclei length (µm) in *D. melanogaster* males expressing different *Mst77F* orthologs. Sperm nuclei from transgenic flies with just one copy of *Mst77F-mel (mel/KO)* are shorter than those with two copies of *Mst77F-mel (mel/mel)*. *Mst77F-sim* homozygous males *(sim/sim)* do not have a significantly different sperm nuclear length than *Mst77F-mel* homozygotes. In contrast, *Mst77F-yak* or *Mst77F-eug* homozygous males *(yak/yak, eug/eug)* have reduced sperm nuclear length, whereas *Mst77F-tak* homozygous males *(tak/tak)* have more extended sperm nuclei. **(B)** Sperm nuclei from *D. yakuba* with no (*KO/KO)* or one copy of *Mst77F-yak (WT-yak/KO)* are shorter than those with two copies of WT-*Mst77F-yak (WT-yak/WT-yak)*. Statistical comparisons were performed using unpaired Student's *t*-tests (*p<0.05; **p<0.01; ****p < 0.0001; ns, not significant).

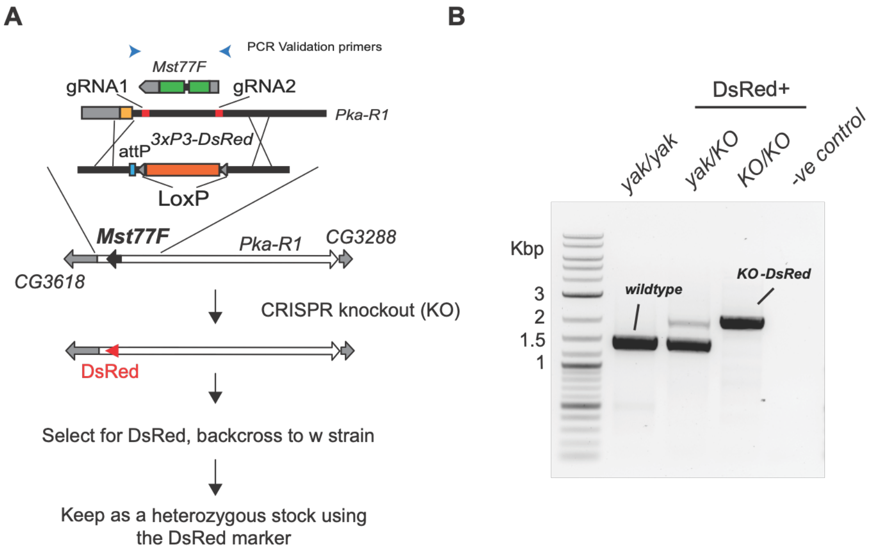


**Fig. S9: Generation and validation of *Mst77F* knockouts in *Drosophila yakuba*. (A)** Schematic illustrating the locations of guide RNA, gRNA1 and gRNA2, and a repair construct carrying a 3×P3-DsRed marker, an attP site and loxP sites. PCR validation primer sites are shown as blue arrowheads. *Mst77F-yak* knockout (*KO*) flies were generated using CRISPR-Cas9 in *D. yakuba*. We replaced the endogenous *Mst77F-yak* gene with *DsRed,* which was subsequently used for selection, strain establishment, and maintenance. Paternal genotype contribution was validated using offspring marker phenotypes *(e.g., DsRed* eye color). **(B)** PCR genotyping of flies carrying the *Mst77F-yak KO* reveals bands corresponding to the wild-type *Mst77F* and *KO* (*DsRed*) alleles in different genotypes, as well as a negative control.

**Data S1. Primer sequences used in this study.**

**Data S2. Progeny counts from all the crosses.**

**Data S3. Quantification of Western blot analyses.**

**Data S4. Quantification of seminal vesicle sizes.**

**Data S5. Sperm counts from all genotypes.**
